# YY1 cistrome analysis uncovers an essential requirement of the YY1:BRD4-PFKP regulatory axis for promoting tumorigenesis of castration-resistant prostate cancer

**DOI:** 10.1101/2020.09.17.302596

**Authors:** Chenxi Xu, Yi-Hsuan Tsai, Phillip Galbo, Weida Gong, Aaron J. Storey, Yuemei Xu, Stephanie D. Byrum, Young E. Whang, Joel S. Parker, Samuel G. Mackintosh, Ricky D. Edmondson, Alan J. Tackett, Jiaoti Huang, Deyou Zheng, H. Shelton Earp, Gang Greg Wang, Ling Cai

## Abstract

Castration-resistant prostate cancer (CRPC) is a terminal disease, demanding a better understanding of its pathogenesis. Targeted therapy needs to be developed for CRPC due to its heterogeneity and resistance to current treatments. Here, through cistrome study of YY1, a transcription factor significantly overexpressed during prostate cancer progression, we identify a YY1-PFKP axis to be essential for CRPC tumorigenesis. Depletion of YY1 in independent CRPC models dramatically reduced tumor cell growth *in vitro* and delayed oncogenic progression *in vivo*. Importantly, YY1 functions as a master regulator of prostate tumor metabolism including the Warburg effect and mitochondria respiration. Loss-of-function and rescue studies further reveals a mechanistic underpinning in which YY1 directly binds and trans-activates *PFKP*, a gene encoding the rate-limiting enzyme for glycolysis, significantly contributing to the YY1-enforced oncogenic phenotypes such as enhanced tumor cell glycolysis and malignant growth. Additionally, a vast majority of gene-regulatory element in advanced prostate cancer cells are bound by YY1, lending a support for its role as a master regulator of prostate cancer progression. YY1 interactome studies point to bromodomain-containing coactivators in prostate cancer, which act as functional partners of YY1 to potentiate YY1-related target gene activation. Altogether, this study unveils an unexplored YY1:BRD4-PFKP oncogenic axis operating in advanced prostate cancer with implications for therapy.

## Introduction

Prostate cancer is the second leading cause of cancer-related death for men in the United States. Standard treatment of prostate cancer with anti-androgen agents fails inevitably due to development of therapy resistance and castration-resistant prostate cancer (CRPC), a terminal disease^1^. Therefore, mechanistic understanding of CRPC pathogenesis, as well as design of novel means to specially target CRPC vulnerabilities, would greatly benefit clinical outcome of the affected patients.

Yin Yang 1 (YY1), a ubiquitously expressed transcription factor, was previously shown to have dual roles in both gene activation and repression^2,3^. It belongs to the GLI-Kruppel zinc finger protein family and carries four conserved C2H2 zinc fingers^4^. As a multifunctional protein, YY1 is involved in various biological and physiological processes including cell proliferation, lineage specification and embryonic development^5^. Emerging evidence indicates that YY1 also play important roles in malignant diseases including cancer. Our analysis of paired normal and tumor patient samples identified YY1 to be significantly overexpressed during prostate cancer progression. However, YY1’s function and regulated cistrome in CRPC are not studied to date.

Aerobic glycolysis, known as the Warburg effect, is essential for cancer to acquire energy and metabolize nutrients for synthesis of macromolecular precursors, in order to sustain high rates of cell proliferation^6^. It has been reported that primary prostate cancers are metabolically different from many other solid tumors, due to their enhanced reliance on oxidative phosphorylation (OXPHOS) in mitochondria, leading to a modest level of glucose uptake^7,8^. Other studies, however, have shown increased glycolysis or the Warburg effect also correlated with disease progression and poor prognosis among the advanced prostate cancers^8^. Thus, the exact molecular mechanism underlying glycolysis regulation in advanced prostate tumors remains elusive, although rising evidence points to possible deregulation of glycolytic enzymes^9^. During glycolysis, phosphofructokinase 1 (PFK1) plays a critical role through catalyzing fructose 6-phosphate (F6P) to fructose 1,6-biphosphate (F1,6BP), one of the rate-limiting steps in glycolysis^10^. PFK1 has 3 isoforms, namely, PFKP (Phosphofructokinase, Platelet), PFKM (Phosphofructokinase, Muscle) and PFKL (Phosphofructokinase, Liver). While all isozymes are expressed in many tissues, PFKP and PFKM are mainly present in platelet and muscle, respectively, whereas PFKL is predominant in liver and kidney^11^. PFKP has recently been also shown to be prevalent in breast cancer, lung cancer and glioblastoma^12–14^. The role of PFKP and its regulation in CRPC remains unexplored.

Here, we report YY1 to be significantly elevated among patient-derived CRPC samples, relative to benign prostate controls, and to be essential for CRPC tumorigenesis in multiple in vitro and in vivo CRPC models. We also use integrative genomics approach with RNA-seq and ChIP-seq to determine the YY1-regulated cistrome in CRPC. Strikingly, we found that YY1 binds a vast majority of gene-regulatory elements (demarcated by H3K27ac) and potentiates various gene-expression programs related to metabolic pathways such as mitochondria respiration and the Warburg effect, thereby profoundly affecting metabolism of advanced prostate tumor cells. A detailed loss-of-function and rescue study points to *PFKP*, a rate-limiting glycolysis enzyme, directly bound and activated by YY1. Oncogenic actions of YY1 is at least partially achieved via PFKP which significantly enhances tumor cell glycolysis, proliferation, soft agar-based colony formation and xenografted tumor growth in mice. Proteomics-based YY1 interactome studies identify bromodomain proteins (BRD2 and BRD4) as YY1’s functional partners, co-mediating transcriptional activation of many metabolic genes including *PFKP*. Taken together, this study shows YY1 as a master regulator of prostate tumorigenesis, unveils a previously unknown oncogenic axis involving YY1:BRD4-PFKP, and elucidates the molecular mechanism underlying altered metabolism of CRPC, implicative of new therapeutic strategies for treatment of lethal CRPCs.

## Results

### YY1 shows significant upregulation among primary samples of prostate cancer patients

YY1 was previously reported to be highly expressed in breast and colorectal cancer^15^. To assess relevance of YY1 in prostate cancer, we first examined the publicly available prostate cancer datasets^16,17^ and found the *YY1* mRNA levels to be significantly elevated in tumors compared to adjacent benign tissues (Fig. 1a-b). Next, we performed immunohistochemical (IHC) staining with thirteen paired tumor and normal tissues from prostate cancer patients. We observed that the YY1 protein level was significantly increased in nuclei of tumors, compared to their respective adjacent benign controls (Fig. 1c-d and Supplementary Fig. 1; *P* = 0.0071). By immunoblots, we further verified upregulation of YY1 in prostate tumor relative to paired benign tissues (Fig. 1e). Furthermore, we performed YY1 IHC staining with tissue microarrays that contained a larger panel of benign prostates and samples representing different stages of prostate tumors, including primary adenocarcinoma and CRPC, and found a significantly higher level of nuclear YY1 in CRPC, compared to normal controls (*P* = 0.0170) and adenocarcinomas (*P* = 0.0339), as demonstrated by representative IHC images (Fig. 1f) and quantitative analysis (Fig. 1g). Overall, these results lend a support for a physiological involvement of YY1 in advanced prostate cancer pathogenesis, including CRPC.

**Figure 1.**
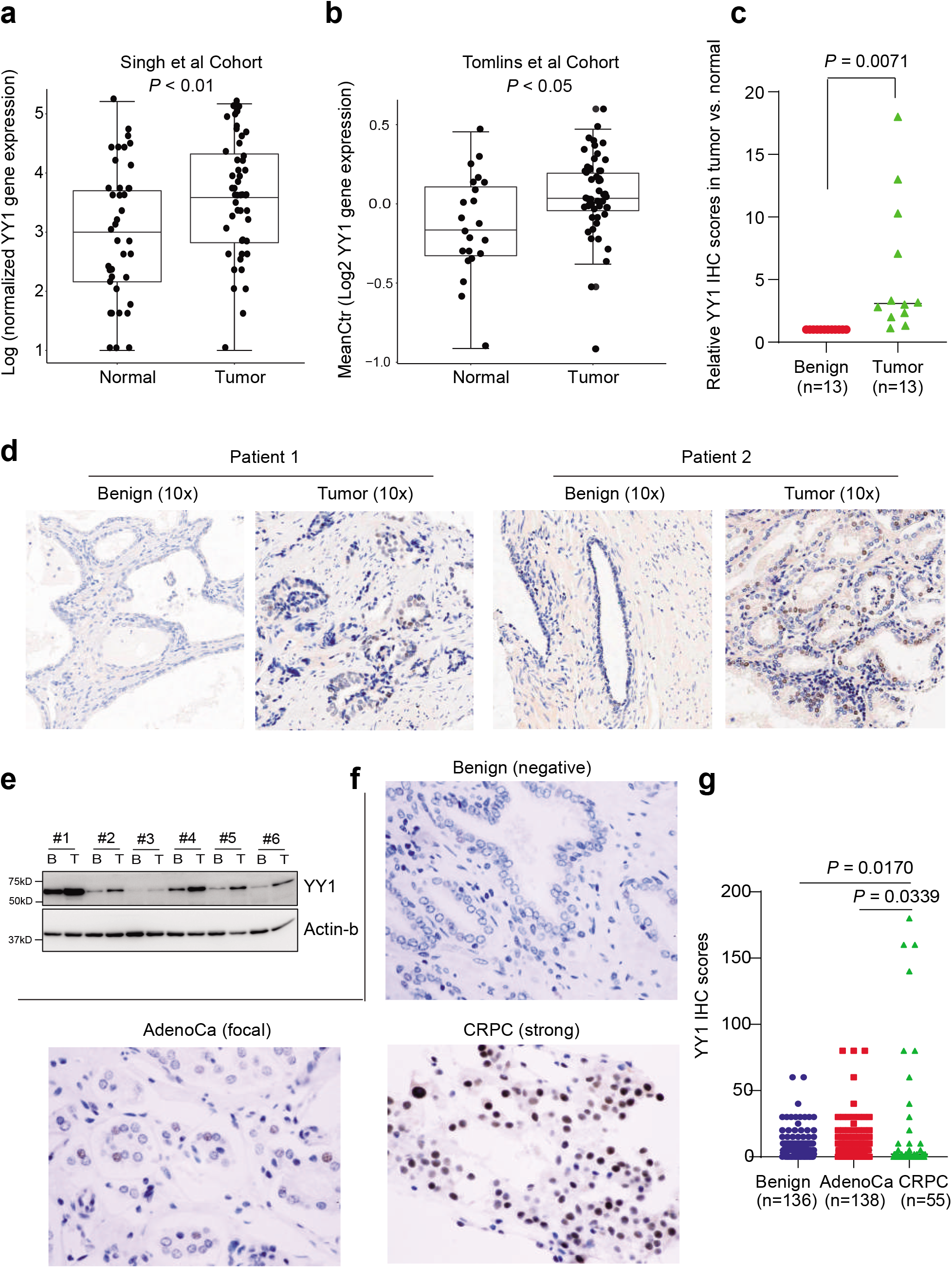
YY1 shows significantly higher expression among samples of prostate cancer patients, when compared to benign tissues. **(a-b)** Boxplots showing overall *YY1* mRNA levels among patient cohorts reported in (Singh et al., 2002) (panel **a**) and (Tomlins et al., 2007) (**b**), relative to their respective normal controls. **(c-d)** Representative images (panel **d**; 10x) and quantification (panel **c**) of YY1 immunohistochemistry (IHC) staining of thirteen paired normal and tumor tissues from prostate cancer patients. **(e)** Immunoblotting for YY1 using total lysates of the paired normal/benign (B) and tumor (T) tissues from prostate cancer patients. β-actin acts as a loading control. **(f-g)** Representative images (panel **f**) and quantification (panel **g**) of YY1 IHC staining by using tissue microarrays (TMA) that contains samples of benign prostates (n=136), primary prostate adenocarcinoma (n=138) and CRPC (n=55).

### YY1 promotes the growth and colony formation of prostate cancer cells *in vitro*, as well as tumorigenesis in xenografted animal models

Next, we sought to determine the role for YY1 in prostate cancer tumorigenesis. Using two independently validated YY1-targeting shRNAs (sh#94 and sh#98), we performed YY1 knockdown (KD) in two androgen-independent CRPC models, 22Rv1 and C4-2 cells (Fig. 2a-f). YY1 KD significantly decreased tumor cell proliferation in liquid culture (Fig. 2b, 2e) and colony formation in soft agar, a surrogate assay of transformation (Fig. 2c, 2f). Similar phenotypes were observed post-KD of YY1 in LNCaP cells, a prostate cancer model showing androgen dependency (Supplementary Fig. 2a-b). Furthermore, re-introduction of an shRNA-resistant YY1 into 22Rv1 cells with endogenous YY1 depleted was able to restore both tumor cell growth and colony formation, ruling out potential off-target effects of the used shRNA (Fig. 2g-i). In accordance with shRNA-mediated YY1 KD, YY1 depletion via two independent sgRNAs through a CRISPR/Cas9 system, or via siRNA, all led to decreased cell proliferation (Fig. 2j-k, Supplementary Fig. 2c-d). To further determine whether YY1 is important for CRPC tumorigenesis *in vivo*, we subcutaneously xenografted the 22Rv1 cells, stably transduced either with control or YY1-targeting shRNAs, into castrated NOD/scid/gamma (NSG) mice. 22Rv1 xenografts in the YY1 KD cohort grew significantly slower relative to controls (Fig. 2l-m). Additionally, we validated YY1 KD in the tumor xenografts (Fig. 2n). Altogether, we conclude that YY1 is crucial for CRPC growth *in vitro* and *in vivo*.

**Figure 2.**
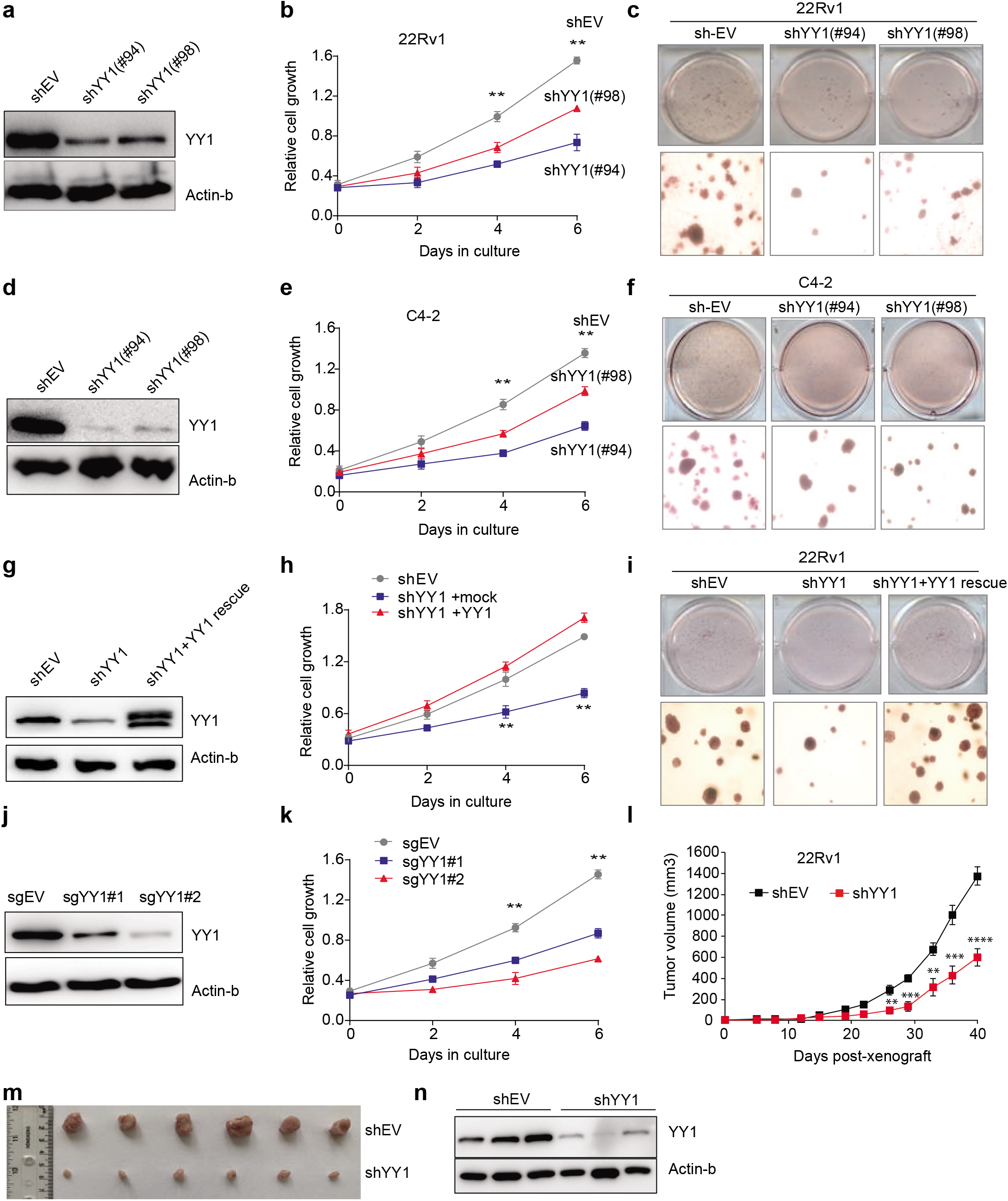
YY1 is required for malignant growth of prostate tumor cells *in vitro* and *in vivo*. **(a-f)** Immunoblotting for YY1 (panels **a** and **d**), measurement of proliferation (**b** and **e**) and assessment of soft agar-based growth (**c** and **f;** with representative images shown) after shRNA-mediated stable knockdown of YY1 (KD by sh#94 or sh#98) in either 22Rv1 (**a-c**) or C4-2 (**d-f**) cells, relative to transduction of empty vector (shEV). \*\**P*<0.01. **(g-i)** Immunoblotting for YY1 (panel **g**), measurement of proliferation (**h**) and assessment of soft agar-based growth (**i;** with representative images shown) post-rescue of YY1 (using an exogenous HA-tagged YY1) in 22Rv1 cells with endogenous YY1 knocked down (by sh#94). \*\**P*<0.01. **(j-k)** Immunoblotting for YY1 (panel **j**) and measurement of proliferation (**k**) after the CRISPR/cas9-mediated knockout of YY1 (sgYY#1 or sgYY#2) in 22Rv1 cells, relative to transduction of empty vector (sgEV). \*\**P*<0.01. **(l-m)** Summary of changes in the tumor volume (panel **l**), following subcutaneous transplantation of stable shEV-(balck) or shYY1-expressing (red) 22Rv1 cells into castrated NSG mice (n = 6 per group). Data presented are mean ± SEM (n = 6 mice for each group). Statistical significance was determined by two-way ANOVA (** *P*< 0.005, *** *P*< 0.0005, **** *P*< 0.0001). The image of representative tumors is shown in **m**. **(n)** Immunoblotting for YY1 in the 22Rv1 tumor xenografts isolated from the indicated NSG cohort (shEV or shYY1).

### YY1 directly binds a set of tumor metabolism-related genes, potentiating their transcription

To gain insight into molecular mechanisms underlying the YY1-mediated CRPC tumorigenesis, we profiled 22Rv1 cell transcriptomes by RNA-seq, which revealed differentially expressed genes (DEGs) caused by YY1 KD (Fig. 3a and Supplementary Table 1). Gene Set Enrichment Analysis (GSEA) showed that genes upregulated by YY1 are enriched in energy metabolism pathways including glycolysis (Fig. 3b, upper panels), oncogenes involved in prostate cancer (Fig. 3b, left/bottom), and, as expected, the YY1 targets (Fig. 3b, right/bottom). Notably, a set of metabolic enzymes involved in glycolysis were downregulated upon YY1 depletion (Fig. 3c). To validate this regulatory function of YY1, we additionally performed RNA-seq post-KD of YY1 in C4-2 cells, another CRPC model, and subsequent GO and GSEA analyses revealed similar striking enrichments of metabolic pathways among genes positively controlled by YY1 (Supplementary Fig. 3a-d, and Supplementary Table 2). Importantly, we identified the DEGs common to both independent CRPC models after YY1 ablation, hereafter termed “the YY1 signature genes in CRPC” (Fig. 3d and Supplementary Table 3), which included a number of metabolism-associated genes such as *PFKP, ALDOC, OGDHL* and *NDUFA4L2*. Using qRT-PCR, we further confirmed downregulation of these metabolic genes upon YY1 loss, relative to control, in both 22Rv1 and C4-2 cells, with the change in *PFKP* expression being most prominent (Fig. 3e-f).

**Figure 3.**
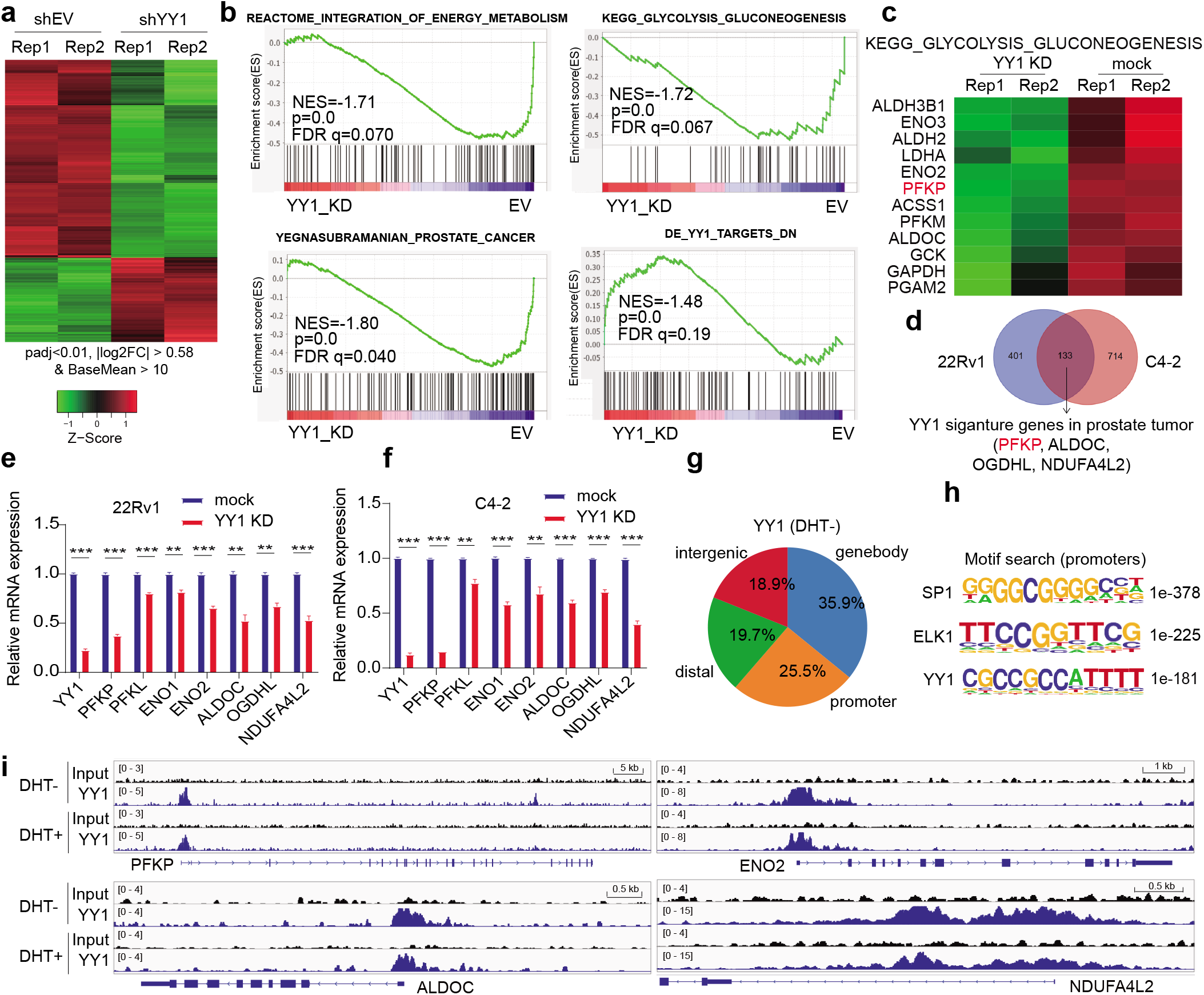
YY1 directly binds to and positively regulates metabolic genes in prostate tumor. **(a)** Heatmap showing the expression levels of differentially expressed genes (DEGs) identified due to YY1 KD relative to mock (shEV) in 22Rv1 cells, with two biological replicates (Rep 1 and 2) per group. Threshold of DEG is set at the adjusted DESeq *P* value (padj) less than 0.01 and fold-change (FC) over 1.5 for transcripts with mean tag counts of at least 10. **(b)** GSEA revealing that, relative to mock, YY1 KD is positively correlated with downregulation of the indicated genesets related to energy metabolism, glycolysis or prostate cancer, and negatively correlated with upregulation of the indicated YY1-repressed geneset (bottom/right) in 22Rv1 cells. **(c)** Heatmap showing expression of the indicated glycolysis-related genes in 22Rv1 cells after YY1 KD, relative to mock. **(d)** Venn diagram showing overlap between the YY1-upregulated genes as identified by RNA-seq in 22Rv1 (left) and C4-2 (right) cells. Threshold of DEG is set at the adjusted DESeq *P* value (padj) less than 0.01 and fold-change (FC) over 1.5 for transcripts with mean tag counts of at least 10. **(e-f)** RT-qPCR of YY1 (far left) and the indicated metabolic gene in 22Rv1 (**e**) and C4-2 (**f**) cells post-KD of YY1, compared with mock. Y-axis shows averaged fold-change ±SD of three independent experiments after normalization to beta-Actin and then to mock-treated. ** *P* <0.01, *** *P* <0.001. **(g)** Pie chart showing genomic distribution of the called YY1 peaks in 22Rv1 cells. **(h)** Motif search analysis revealing the most enriched motifs at the called YY1 peaks found at gene promoters. **(i)** IGV browser views of chromatin input and YY1 ChIP-seq peaks (depth normalized) at the indicated metabolic pathway genes in 22Rv1 cells, ligand-starved followed by treatment with vehicle (DHT-) or dihydrotestosterone (DHT+).

We next performed ChIP-seq to determine genome-wide YY1 binding sites in 22Rv1 cells that were ligand-starved followed by treatment with vehicle (DHT-) or AR agonist (DHT+) (Supplementary Fig. 3e). YY1 binding patterns are highly similar between vehicle- and DHT-treated cells (data not shown); thus, we chose the YY1 ChIP-seq peaks called from the vehicle-treated cells for further analysis, as this condition more closely resembles CRPC. Genomic localization analysis showed approximately 25% of the YY1 peaks at promoters, ~36% in the gene body and the rest (~39%) at putative intergenic and distal enhancers (Fig. 3g). As expected, YY1 motif was among the most enriched motifs identified within the YY1 peaks (Fig. 3h). Also, YY1 peaks were found at almost all of the YY1-upregulated genes defined by RNA-seq, including metabolic genes *PFKP, ALDOC, ENO2* and *NDUFA4L2* (Fig. 3i). ChIP-qPCR of YY1 further validated its strong enrichment at metabolic gene promoters (Supplementary Fig. 3f). Taken together, our genome-wide-profiling integration analyses lend a strong support for a direct involvement of YY1 in upregulation of genes related to cell metabolism in prostate cancer.

### YY1 potentiates prostate tumor cell glycolysis via PFKP

Next, we sought to assess whether YY1 regulates prostate cancer cell metabolism and measured oxygen consumption rate (OCR) in both 22Rv1 and C4-2 cells post-KD of YY1 versus control. Indeed, YY1 depletion led to the significantly reduced levels of basal OCR and maximal respiratory capacity (Fig. 4a-b). To explore how YY1 regulates the Warburg effect of prostate cancer, we further measured the basal extracellular acidification rate (ECAR), a key marker of glycolysis, and observed it to be dramatically decreased after YY1 loss in three independent prostate tumor models, 22Rv1, C4-2 and LNCaP cells (Fig. 4c-d, Supplementary Fig. 4a). In agreement with these metabolic changes caused by YY1 loss, overexpression of wildtype (WT) YY1, but not its mutant form with the N-terminal domain deleted, which was previously reported to be transactivation-defective^18^, was able to further enhance glycolysis of 22Rv1 and C4-2 cells (Fig. 4e–4g and Supplementary Fig. 4b). Such a requirement of YY1’s transactivation domain for promoting tumor cell glycolysis is in line with a role for YY1 in transactivation of glycolysis-related genes revealed by RNA-seq.

**Figure 4.**
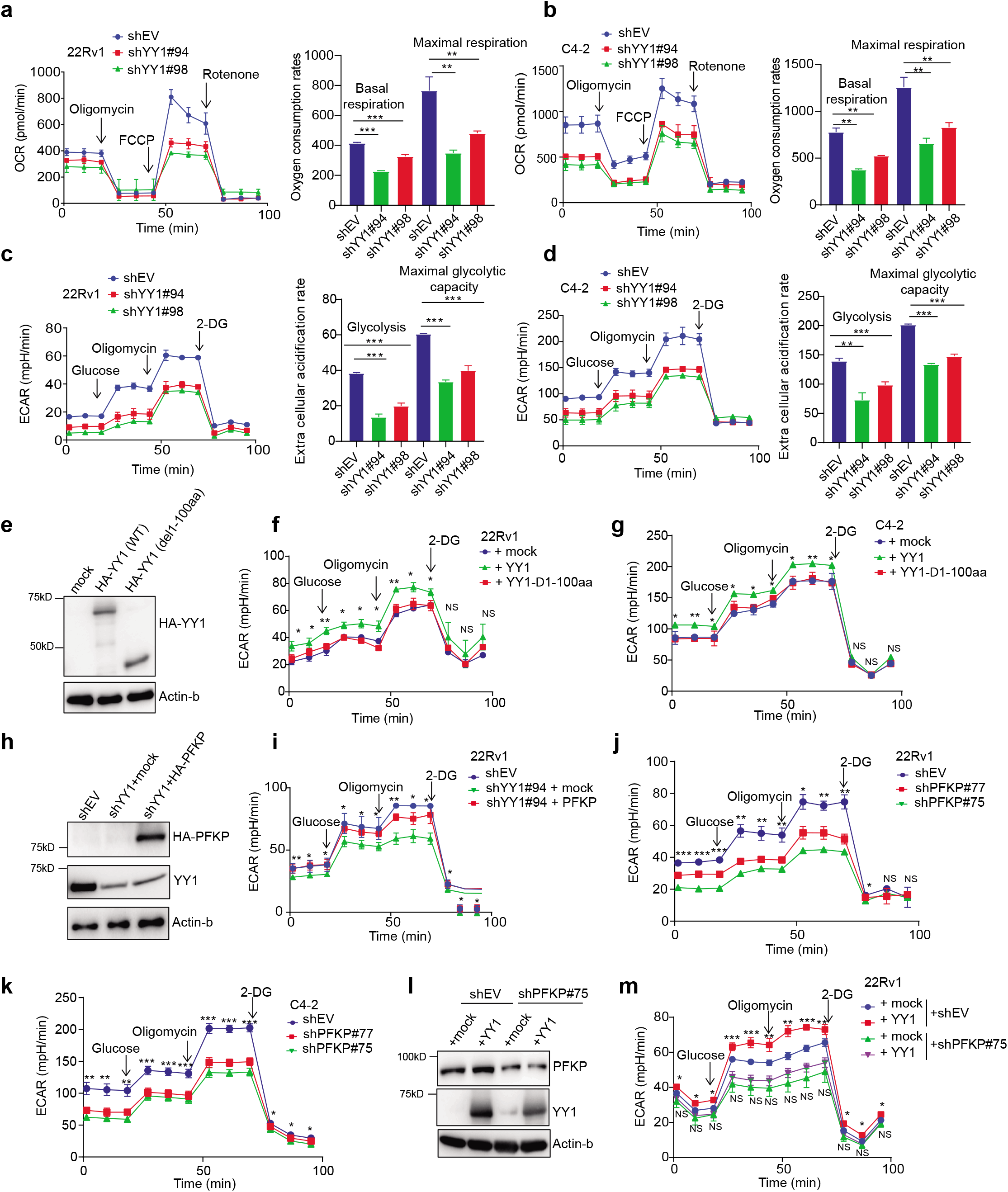
YY1 potentiates prostate tumor cell glycolysis via PFKP. **(a-b)** Measurement of oxygen consumption rate (OCR) and mitochondrial bioenergetics in 22Rv1 (panel **a**) and C4-2 (**b**) cells post-KD of YY1, compared to mock (shEV), by using the Seahorse XF-24 extracellular flux analyzer. Injection of compounds during the assay is highlighted in left panels. Data quantifications are shown as mean±SEM in right panels. ** *P*<0.01, *** *P* <0.001. **(c-d)** Measurement of extra cellular acidification rate (ECAR) in 22Rv1 (panel **c**) and C4-2 cells (**d**) post-KD of YY1, compared to mock (shEV). Injection of compounds during the assay is highlighted in left panels. Data quantifications are shown as mean±SEM in right panels. ** *P*<0.01, *** *P* <0.001. **(e-g)** Immunoblotting for YY1 (panel **e**; anti-HA immunoblots in 22Rv1 cells), as well as ECAR measurements in 22Rv1 (**f**) or C4-2 (**g**) cells post-transduction of the indicated HA-tagged YY1, either WT or with a N-terminal transactivation domain deleted (Δ1-100aa). * *P*<0.05; ** *P* <0.01; NS, not significant. **(h-i)** Endogenous YY1 and anti-HA immunoblotting (panel **h**) and ECAR measurements (**i**) post-transduction of an HA-tagged PFKP (**h**, lane 3), relative to mock (**h**, lane 2), into the YY1-depleted 22Rv1 cells (lanes 2-3). The shEV-transduced cells (lane 1) serve as control. * *P*<0.05; ** *P* <0.01; NS, not significant. (**j-k**) ECAR measurements in 22Rv1 (panel **j**) or C4-2 (**k**) cells post-KD of PFKP (sh#75 or sh#77), compared to mock (shEV). * *P*<0.05; ** *P* <0.01; *** *P* <0.001. (**l-m**) YY1 and PFKP immunoblotting (panel **l**) and ECAR measurements (**m**) post-transduction of an HA-tagged YY1 (**l**, lanes 2 and 4), relative to vector mock (**l**, lanes 1 and 3), into 22Rv1 cells with stable expression of either shEV (lanes 1-2) or YY1-targeting shRNA (lanes 3-4). * *P*<0.05; ** *P* <0.01; NS, not significant.

Next, we aimed to determine which YY1’s downstream metabolic gene target(s) mediates glycolysis in prostate tumor. Among the commonly YY1-upregulated signature transcripts *PFKP, ALDOC* and *ENO2* (Fig. 3c), we found that overexpression of ENO2 or ALDOC failed to significantly rescue the glycolysis defects caused by YY1 depletion (Supplementary Fig. 4c-f). In contrast, the restored expression of PFKP largely rescued YY1 loss-related defects (Fig. 4h-i). Consistent with this finding, KD of PFKP using either of two independently validated hairpins (sh#75 and sh#77) significantly diminished the rate of glycolysis in both 22Rv1 and C4-2 cells (Fig. 4j-k); likewise, depletion of PFKP almost completely abolished the increased glycolysis caused by YY1 overexpression (Fig. 4l-m).

To this end, we show that YY1 exerts an essential role in potentiating prostate cancer cell glycolysis, an event heavily relying on upregulation of *PFKP* expression.

### PFKP is a direct onco-target of YY1 in prostate cancer

We next examined how YY1 regulates PFKP expression. First, the Volcano plots based on RNA-seq profiles of 22Rv1 (Fig. 5a) and C4-2 (Fig. 5b) cells both pointed to *PFKP* among the most altered transcripts upon YY1 depletion. Consistently, YY1 ablation in these CRPC cells led to a significant decrease of PFKP protein levels, relative to control (Fig. 5c). Meanwhile, rescue of YY1 loss by an exogenous YY1^WT^ restored the PFKP protein level in these cells (Fig. 5d, middle vs left lanes), an effect not seen with YY1^S365D^, a DNA-binding-defective mutant^19^ (Fig. 5d, right lanes). This indicates that *PFKP* induction by YY1 is DNA-binding-dependent, in agreement with our YY1 ChIP-seq results showing a direct binding of YY1 to the *PFKP* promoter (Fig. 5e). Indeed, YY1^WT^, but not its YY1^S365D^ mutant, increased transcription activity from a luciferase-based reporter that carried either a 2Kb-long^20^ (Fig. 5f) or 575bp-long region upstream of PFKP’s transcriptional start site (TSS; Fig. 5g; based on our ChIP-seq data in Fig. 5e). Furthermore, there are four YY1 core binding motifs, CCAT^21^, within the YY1 binding peaks at the *PFKP* promoter (i.e., −575 to +37 bp off TSS; Supplementary Fig. 5). Systematic mutagenesis of these four YY1-binding motifs (Fig. 5h; mutation of CCAT to CGGT) revealed the first and third CCAT motifs, but not the second and fourth ones, to be essential for the *PFKP* promoter-driven transactivation activity in both 22Rv1 and C4-2 cells (Fig. 5i-j). Additionally, relative to the single motif mutation, compound mutation of the first and third YY1 motifs further decreased the *PFKP* promoter activity (Fig. 5i-j; last panels). Thus, YY1 transactivates *PFKP* through directly binding its promoter, particularly through the first and third conserved CCAT motifs.

**Figure 5.**
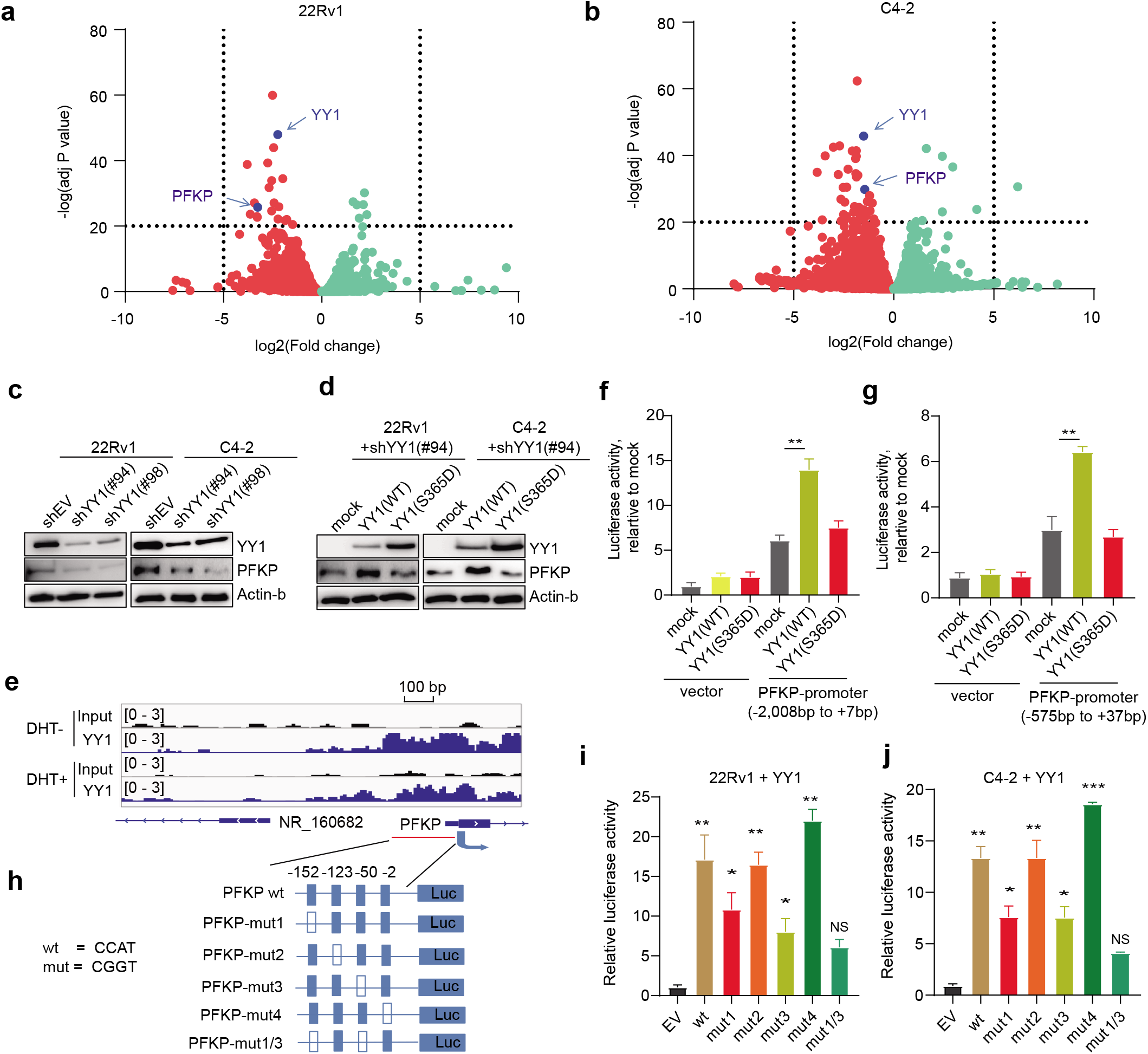
PFKP is a direct onco-target of YY1 in prostate cancer. **(a-b)** Volcano plots based on RNA-seq profiles highlight *PFKP* among the most altered transcripts upon YY1 depletion in 22Rv1 (**a**) and C4-2 (**b**) cells. **(c)** YY1 and PFKP immunoblots post-KD of YY1 in 22Rv1 (left) and C4-2 (right) cells. **(d)** YY1 and PFKP immunoblots post-transduction of an HA-tagged YY1^WT^ or YY1^S365D^, a DNA-binding-defective mutant, into 22Rv1 (left) and C4-2 (right) cells. **(e)** IGV views of the YY1 ChIP-seq profile on the proximal promoter region of *PFKP*. **(f-g)** Relative luciferase activities from a reporter that carries promoter region of *PFKP*, either from −2,008 to +7 (**f**) or −575 to +37 (**g**), in 22Rv1 cells expressed with HA-tagged YY1^WT^ or YY1^S365D^. Y-axis shows averaged fold-change ±SD of three independent experiments after normalization to internal luciferase controls and then to mock samples (EV-expressing). ** *P*<0.01. **(h)** Scheme showing mutations (mut) of putative YY1-binding sites within the *PFKP* promoter. Empty box denotes the mutated YY1-binding site. **(i-j)** Relative luciferase activities from a PFKP promoter reporter, either WT or that carrying the indicated mutation of putative YY1-binding sites (refer to panel **h**), following overexpression of YY1^WT^ into either 22Rv1 (**i**) or C4-2 cells (**j**). Y-axis shows averaged fold-change ±SD of three independent experiments after normalization to internal luciferase controls and then to EV-expressing control samples (far left). * *P* <0.05; ** *P* <0.01; *** *P* <0.001.

### PFKP is critically involved in prostate cancer tumorigenesis *in vitro* and *in vivo*

Since PFKP functions as a key rate-limiting enzyme in glycolysis, we next examined its involvement in prostate cancer tumorigenesis, which was previously unexplored. First, examination of *PFKP* across different transcriptome datasets of prostate cancer samples^17,22^ showed its significantly higher expression in prostate tumors, compared to benign tissues (Fig. 6a-b). To further determine the role for PFKP in prostate oncogenesis, we depleted PFKP in both 22Rv1 and C4-2 cells (Supplementary Fig. 6a-b) and observed the significantly decreased levels of *in vitro* proliferation (Fig. 6c) and soft agar-based growth (Fig. 6d), relative to mock. Importantly, overexpression of PFKP in the YY1-depleted 22Rv1 and C4-2 cells partially but significantly rescued the ameliorated proliferation phenotype caused by YY1 loss (Fig. 6e), consistent to PFKP’s rescue effects in the tumor cell metabolic assays (Fig. 4i). In the 22Rv1 cell xenografted mouse model, PFKP loss also dramatically decreased tumor growth *in vivo* (Fig. 6f-h). Thus, PFKP, a downstream target of YY1, is essential for prostate cancer progression.

**Figure 6.**
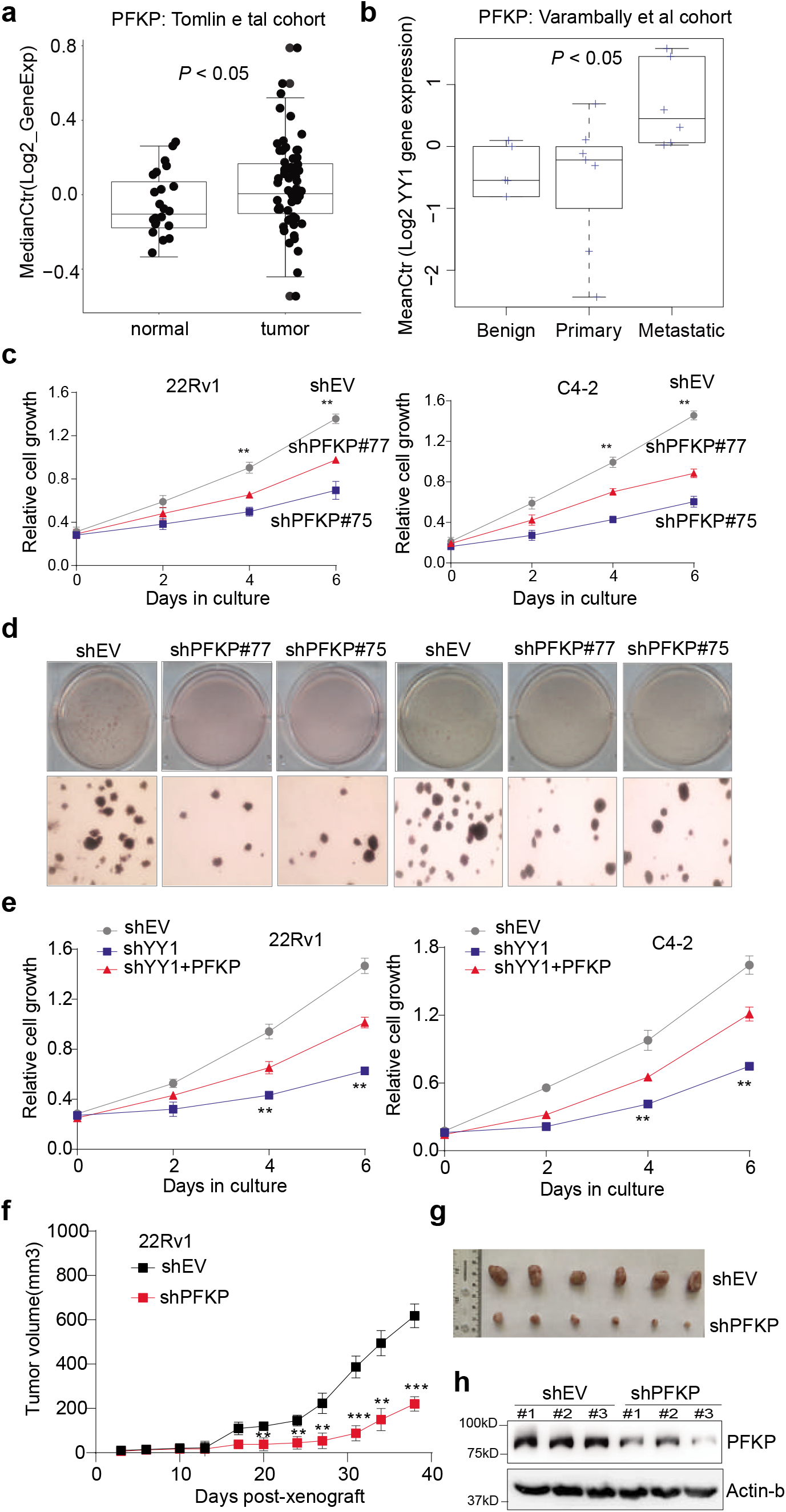
PFKP is critically involved in prostate cancer tumorigenesis *in vitro* and *in vivo*. **(a-b)** Box plots showing the *PFKP* expression levels among samples from the indicated patient cohorts reported in (Tomlins et al., 2007) (panel **a**) or (Varambally et al., 2008) (**b**). **(c-d)** Assays for in vitro proliferation (**c**) and soft agar-based growth (**d**) after PFKP depletion (sh#75 or sh#77), compared to control (shEV), in 22Rv1 (left) and C4-2 cells (right). * *P* <0.05; ** *P* <0.01; *** *P* <0.001. **(e)** Assays for proliferation post-transduction of an exogenous PFKP (red), relative to mock (blue), into the YY1-depleted 22Rv1 (left) and C4-2 cells (right), with non-depleted cells acting as controls (shEV; gray). ** *P* <0.01. **(f-g)** Summary of changes in the tumor volume (**f**), following subcutaneous transplantation of stable shEV-(balck) or shPFKP-expressing (red) 22Rv1 cells into castrated NSG mice (n = 6 per group). Statistical significance was determined by two-way ANOVA (** *P*< 0.005, *** *P*< 0.0005). The image of representative tumors is shown in **g**. **(h)** Immunoblotting for PFKP in the 22Rv1 tumor xenografts isolated from the indicated NSG cohort (shEV or shPFKP).

### Bromodomain-containing protein acts as cofactor of YY1, critically contributing to the YY1-related transactivation of metabolic genes in prostate cancer

To gain further insight into the gene-activation mechanism underlying the YY1-mediated CRPC tumorigenesis, we examined into the YY1 interactome by employing a proximity labeling-based BioID approach^23,24^ (Supplementary Fig. 7a), and a subsequent mass spectrometry-based proteomics analyses identified INO80, YY2 and BRD2 among the most significantly enriched hits as YY1-asscociated factors (Fig. 7a). INO80, a known YY1-interacting proteins^25^, was identified, which validated our method. By co-immunoprecipitation (CoIP), we further verified the interaction between YY1 and BRD2 or a BRD2-related bromodomain protein BRD4 in 22Rv1 cells (Fig. 7b) and C4-2 cells (Supplementary Fig. 7b). We further mapped that the bromodomains of BRD4, which are known to be highly conserved among all BRD family members, were required for interaction with YY1 (Fig. 7c). Acetylated histones such as H3K27ac is known to provide a platform for tethering the BRD4-pTEFb complexes, which in turn boost the release of RNA Pol-II into a productive elongation phase^26^. In agreement, we found a striking overlap between YY1 ChIP-seq peaks with those of H3K27ac in 22Rv1 cells^27^—about 90% of H3K27ac ChIP-seq peaks are co-localized with YY1 (Fig. 7d). Consistently, BRD4 ChIP-seq in 22Rv1 cells revealed significant binding at the YY1 promoter peaks, nearly as strong as YY1, and at most of the YY1 non-promoter peaks (Fig. 7e), as exemplified by those at metabolic genes such as *PFKP* and *ALDOC* (Fig. 7f and Supplementary Fig. 7c). In addition, treatment of 22Rv1 cells with JQ1, an inhibitor of bromodomain proteins BRD2 and BRD4, dramatically decreased overall expression of our defined YY1 signature genes including the metabolism-related ones (Fig. 7g). Using RT-qPCR, we further validated the inhibitory effect of JQ1 on metabolic genes such as *PFKP* (Fig. 7h). Collectively, the YY1-mediated potentiation of a metabolic gene-expression program is enhanced by its coactivators, bromodomain-containing proteins (BRD2/4), in prostate cancer (see a model in Fig. 7i).

**Figure 7.**
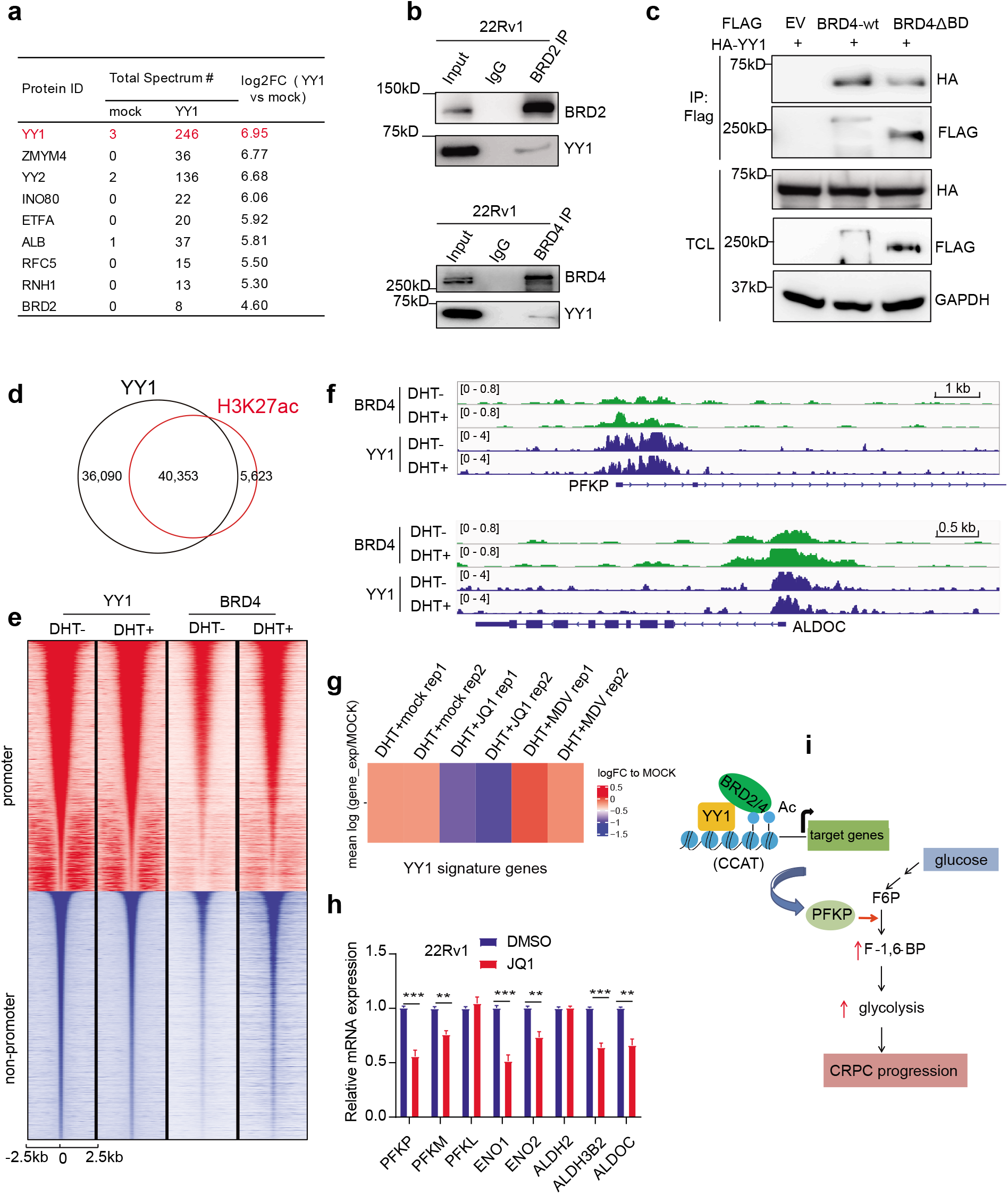
Bromodomain-containing protein acts as co-activator of YY1, potentiating expression of glycolysis-related genes in prostate cancer cells. **(a)** Summary of the top hits identified using BioID and 22Rv1 cells expressing YY1 with a biotin ligase fused to its N-terminus, compared to control cells (EV). **(b)** Co-immunoprecipitation (CoIP) for interaction between endogenous YY1 and bromodomain-containing proteins, BRD2 (top) and BRD4 (bottom), in 22Rv1 cells. **(c)** CoIP for interaction between the exogenously expressed BRD4 (Flag-tagged), either WT or BD domains deleted (ΔBD), and HA-tagged YY1 in 293 cells. TCL, total cell lysate. **(d)** Venn diagram showing overlap between the called YY1 and H3K27ac ChIP-seq peaks identified from the ligand-stripped 22Rv1 cells. **(e)** Heatmap of the YY1 and BRD4 ChIP-Seq read densities at promoter (top) and non-promoter (bottom) YY1 peaks in vehicle-(DHT-) or DHT-treated 22Rv1 cells. The YY1 peaks from vehicle and DHT were combined, sorted, and used to compute ChIP-seq read densities within 5 kb of the YY1 peak centers. **(f)** IGV views of the YY1 and BRD4 ChIP-seq profiles at the indicated glycolytic genes in vehicle-(DHT-) or DHT-treated 22Rv1 cells. **(g)** Heatmap showing overall expression changes fo the YY1 gene signature (defined in Figure 3C) in 22Rv1 cells after drug treatment, either DHT alone, DHT plus JQ1, or DHT plus AR antagonist MDV3100 (MDV). Color bar, mean of the log2FC compared to mock. **(h)** RT-qPCR of the indicated metabolic gene in 22Rv1 post-treatment of DMSO or JQ1 for 8 hours. Y-axis shows averaged fold-change ±SD of three independent experiments after normalization to *beta-Actin* and then to mock-treated. ** *P* <0.01, *** *P* <0.001. **(i)** A model illustrating the role for YY1:BRD4 in potentiating energy metabolism in advanced prostate tumor.

## Discussion

Metabolic reprogramming towards aerobic glycolysis seen in cancer, initially discovered by Otto Warburg, is now appreciated to be a hallmark of tumor, especially advanced ones, to gain survival and growth advantages. Unlike many other solid cancer types, prostate cancer exhibits a unique context- and stage-dependent alteration in metabolism^7^. It is generally viewed that, at its early stage, prostate cancer relies on the increased oxygen consumption, as well as aerobic glycolysis, to support enhanced cell proliferation; the Warburg effect becomes increasingly more pronounced during progression into aggressive, late-stage prostate tumor^8^. However, the molecular underpinning of metabolic reprogramming seen in advanced prostate tumor remains unclear.

Although YY1 was previously reported to interact with AR and regulate expression of prostate tumor marker genes such as *PSA*^28^, there is a general lack of understanding of the role for YY1 in prostate oncogenesis, especially as prostate cancer advances to CRPC. In this report, we show that YY1 plays a pivotal role in prostate cancer progression across independent tumor models such as AR-dependent and CRPC cancer cell lines and in vivo xenografted animal models. Furthermore, we demonstrate that, via direct binding and transactivation of downstream metabolic genes, YY1 is paramount for potentiating metabolism-related programs including mitochondria respiration and glycolysis in these prostate cancer models. This finding is in agreement with what was reported in other biological contexts— for example, skeletal muscle-specific knock-out of YY1 in mice exhibited defective mitochondria morphology and functions^29^; YY1 also activates mitochondria bioenergetics-related genes in B cells^30^ and alters tumor cell metabolism in colon cancer by activating G6PD and the pentose phosphate pathway^31^. In this study, we demonstrated that PFKP, a key metabolic gene whose expression is profoundly affected by YY1 via a direct promoter binding, is essential for YY1-mediated tumor cell glycolysis. Altogether, our findings unveil a previously unexplored axis involving YY1-PFKP, which acts in prostate cancer to sustain aggressive cancer phenotypes. We also favor a view that YY1 likely acts as a master regulator of cancer cell metabolism in many tumor types for sustaining their needs in energy and metabolism controls (through directly activating mitochondrial and/or glycolytic pathways) during progression into advanced diseases. Of note, YY1 was reported to mediate tumorigenesis such as breast cancer and melanoma as well, indicating a multifaceted role of YY1 in oncogenesis^32–35^. Additionally, our study sheds light on potential strategies of targeting metabolism alterations in prostate cancer. By Biotin followed by mass spectrometry, we identify BRD4 family proteins (BRD2 and BRD4) as functional partners of YY1, providing a mechanistic explanation for the YY1-induced transcriptional potentiation of metabolic programs seen in advanced CRPC. Importantly, this pathway is druggable by the bromodomain inhibitors implicating a promising therapy of CRPC.

Equally intriguing is that, besides a transcriptional regulator via direct binding to consensus CCAT motifs (such as that seen at the YY1-targeted PFKP promoter), YY1 was recently shown to function as a three-dimensional genome ‘organizer’ through interactions with additional factors (such as CTCF) for mediating formation of looping between enhancers and promoters^36,37^. How YY1 regulates 3D genome structure of prostate cancer remains an open area for future studies but the current data implicates a direct role in an important node in altering CRPC metabolism. CRPC is a heterogeneous and lethal disease among men with limited therapeutic options, and the gained understanding of CRPC pathogenesis due to this work shall aid in its improved targeted therapies.

## Materials and Methods

### Analysis of public prostate cancer datasets

Public gene expression datasets are from the Singh et al. (2002) study (GSE68907), the Tomlins et al. (2007) study (GSE6099), the Varambally et al. (2008) study (GSE3325). Gene expression data available for the gene of interest were extracted, log2 transformed for each sample. These summarized values were tested for association with sample type (such as benign, primary or metastatic) by ANOVA.

### Tissue Microarray (TMA)

TMAs, produced by the Duke Pathology department, were subjected to YY1 immunohistochemistry (IHC) staining and evaluated in a blinded fashion by the pathologists. Scoring was assessed on the basis of staining intensity from 0 (no staining) to 3 (strong) and percentage of tumor cell expression (1 to 100%), creating a composite score from 0 to 300.

### RNA-seq

RNA was prepared as described before^38^, using 2 million of the 22Rv1 or C4-2 cells stably transduced with shRNAs. Then, complementary DNA was generated, amplified and subjected for library construction using TruSeq RNA Library Preparation Kit v2 (Illumina; catalog# RS-122-2002). The multiplexed RNA-Seq libraries were subject to deep sequencing using the Illumina Hi-Seq 2000/2500 platform according to the manufacturer’s instructions.

### ChIP-seq

YY1 ChIP-seq was carried out as before^38^. Briefly, 22Rv1 cells were first cultured under ligand-starved conditions for three days, followed by a 6-hour drug treatment with vehicle or 10nM of DHT. Cells were cross-linked with 1% formaldehyde at room temperature for 10 minutes, followed by addition of glycine to stop crosslinking. After washing, lysis and sonication, cell chromatin fractions were incubated with antibody-conjugated Dynabeads (Invitrogen) overnight at 4 degree. Chromatin bound beads were then subject to extensive washing and elution. Eluted chromatin was de-crosslinked overnight at 65 degree, followed by protein digestion with proteinase K and DNA purification with QIAGEN PCR purification kit. The obtained ChIP DNA samples were submitted to the UNC-Chapel Hill High-Throughput Sequencing Facility (HTSF) for preparation of multiplexed libraries and deep sequencing with an Illumina High-Seq 2000/2500 platform according to the manufacturer’s instructions. H3K27ac and BRD4 ChIP-seq datasets in 22Rv1 cells were downloaded from the ENCODE project (ENCODE Project Consortium, 2011) and published paper^38^, respectively.

### In vivo tumor growth in xenograft models

All animal experiments were approved by and performed in accord with the guidelines of the Institutional Animal Care and Use Committee (IACUC) at UNC. One million of 22Rv1 cells with stable transduction of shRNA or control empty vector were suspended in 100ul of PBS with 50% Matrigel (BD Biosciences) and then subcutaneously (s.c.) injected into the dorsal flanks of castrated NOD/SCID/gamma-null (NSG) mice bilaterally (carried out by the Animal Studies Core, UNC at Chapel Hill Cancer Center). Tumor growth was monitored twice a week and the tumor volume calculated.

### Statistical analysis

Unless specifically indicated, the unpaired two-tail Student’s t test was used for experiments comparing two sets of data. Data represent mean ± SEM from three independent experiments. *, **, and *** denote *P* values < 0.05, 0.01, and 0.001, respectively. NS denotes not significant.

### Data Availability

RNA-seq and ChIP-seq reads have been deposited to Gene Expression Omnibus (GEO) under accession number GSE153640. The matched input data of 22Rv1 cells (both EtOH and DHT-treated) were previously published by us (available from NCBI GEO #GSE94013) and used herein for normalization of 22Rv1 YY1 ChIP-seq datasets.

## Acknowledgments

We thank UNC’s facilities, including Imaging Core, High-throughput Sequencing Facility (HTSF), Bioinformatics Core, Tissue Procurement Facility, Translational Pathology Laboratory, Tissue Culture Facility and Animal Studies Core for their professional assistance of this work. We graciously thank Drs. Kyung-Sup Kim and Hyun Sil Kim for providing reagents and the Cai, Wang and Earp laboratory members for discussion and technical support. We also acknowledge proteomics support from the University of Arkansas for Medical Sciences Proteomics Core Facility, the IDeA National Resource for Proteomics, the Translational Research Institute through the National Center for Advancing Translational Sciences of the National Institutes of Health, and the Center for Translational Pediatric Research Bioinformatics Core Resource with support through TL1TR003109, P20GM121293, P20GM103625, P20GM103429, S10OD018445, UL1TR003107, and R01CA236209. This work was supported in part by UNC Lineberger Cancer Center UCRF Stimulus Initiative Grant (to L.C.). The cores affiliated to UNC Cancer Center are supported in part by the UNC Lineberger Comprehensive Cancer Center Core Support Grant P30-CA016086. G.G.W. is an American Cancer Society (ACS) Research Scholar and a Leukemia and Lymphoma Society (LLS) Scholar.

## Declaration of interests

J.H. is a consultant for or owns shares in the following companies: Kingmed, MoreHealth, OptraScan, Genetron, Omnitura, Vetonco, York Biotechnology, Genecode and Sisu Pharma. G.G.W. is an inventor of a patent application filed by University of North Carolina at Chapel Hill. G.G.W. received research fund from the Deerfield Management/Pinnacle Hill Company.

## Author contributions

C.X. performed most of the experiments. H.S.E., G.G.W. and L.C. conceived the project, organized and led the study. Y.-H.T. and W.G. conducted analysis of RNA-seq and public cancer datasets under the supervision of J.S.P., G.G.W. and L.C., A.J.S., S.G.M., R.D.E., and S.D.B. performed proteomics analyses using mass spectrometry. A.J.T. supervised the proteomics analysis. P.G analyzed ChIP-seq data under the supervision of D.Z., G.G.W. and L.C. Y.X. and J.H. provided critical guidance and helps on IHC and TMA. C.X., Y.W., J.S.P., J.H., D.Z., H.S.E., G.G.W. and L.C. interpreted the data. L.C. conceived the idea, supervised the work and designed the research. C.X., G.G.W. and L.C. wrote the manuscript with input from other authors.

