## Supplementary materials for "YY1 cistrome analysis uncovers an essential requirement of the YY1:BRD4-PFKP regulatory axis for promoting tumorigenesis of castration-resistant prostate cancer"

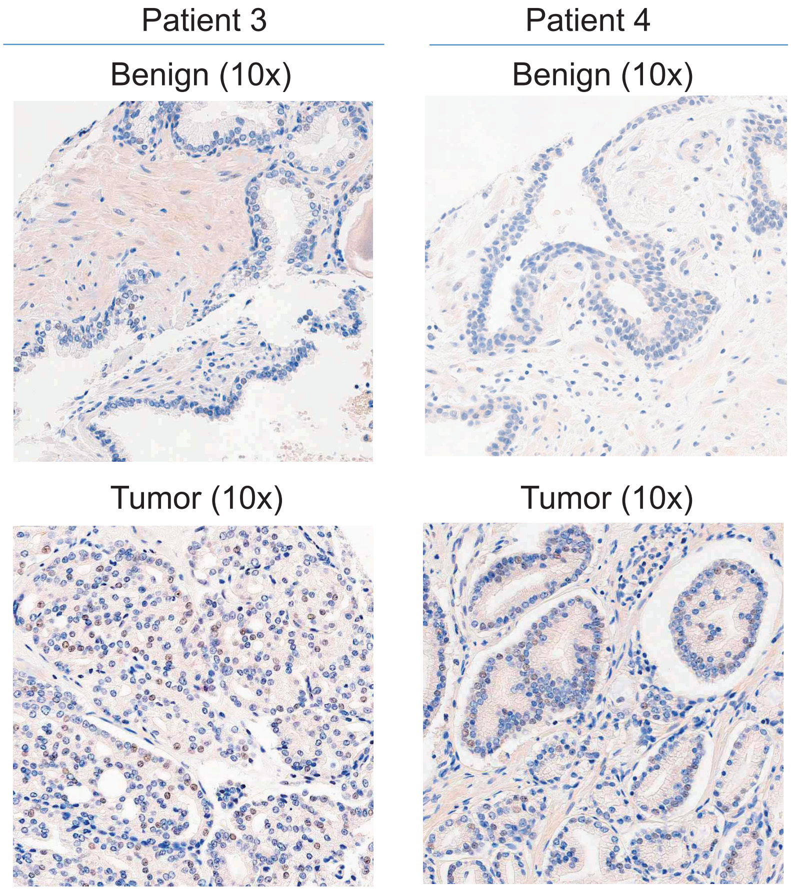


**Supplementary Fig.1: YY1 shows significantly higher expression among samples of prostate cancer patients, when compared to benign tissues.** Representative images of YY1 immunohistochemistry (IHC; 10x) staining of paired normal/benign and tumor tissues from prostate cancer patients.

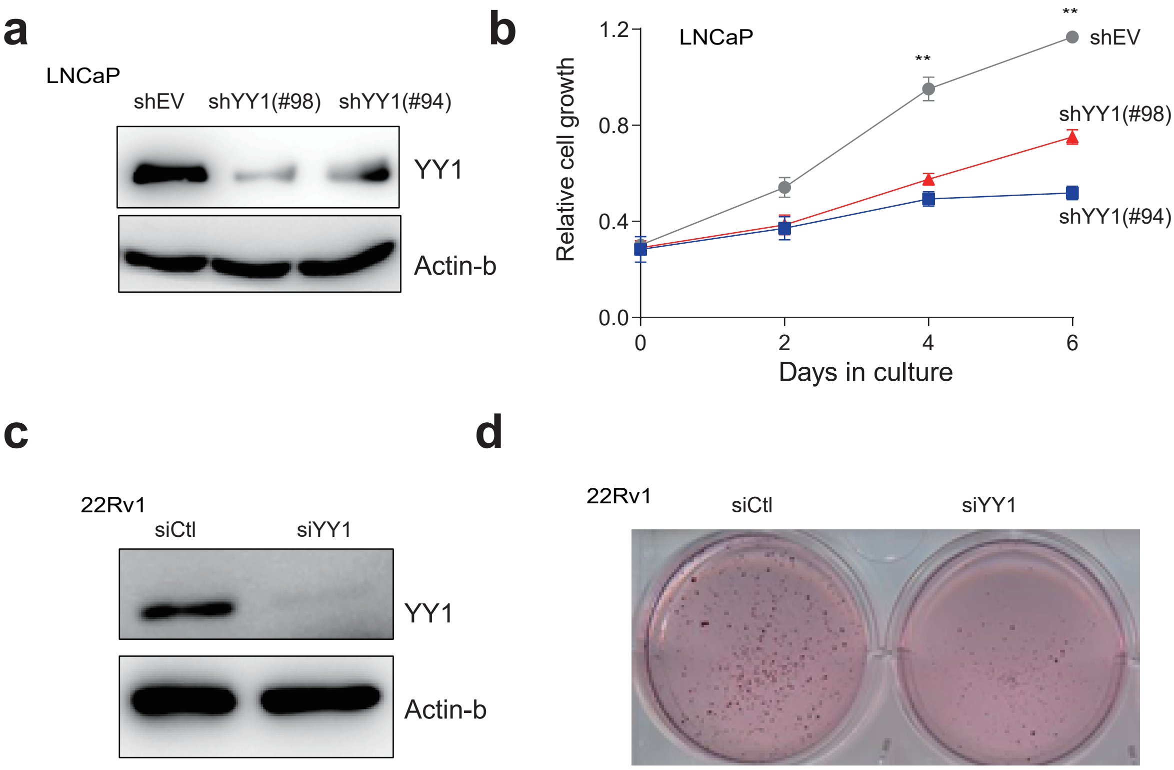


**Supplementary Fig.2: YY1 is required for malignant growth of prostate tumor cells in vitro and in vivo. (a-b)** Immunoblotting for YY1 (panel a) and measurement of proliferation **(b)** after the shRNA-mediated stable knockdown of YY1 (KD by sh#94 or sh#98) in LNCaP cells, relative to transduction of empty vector (shEV). **P<0.01. **(c-d)** Immunoblotting for YY1 (panel **c**) and assessment of the soft agar-based growth (**d**; with representative image shown) post-KD of YY1 via siRNAs (siYY1) in 22Rv1 cells, relative to scramble RNAi controls (siCtl).

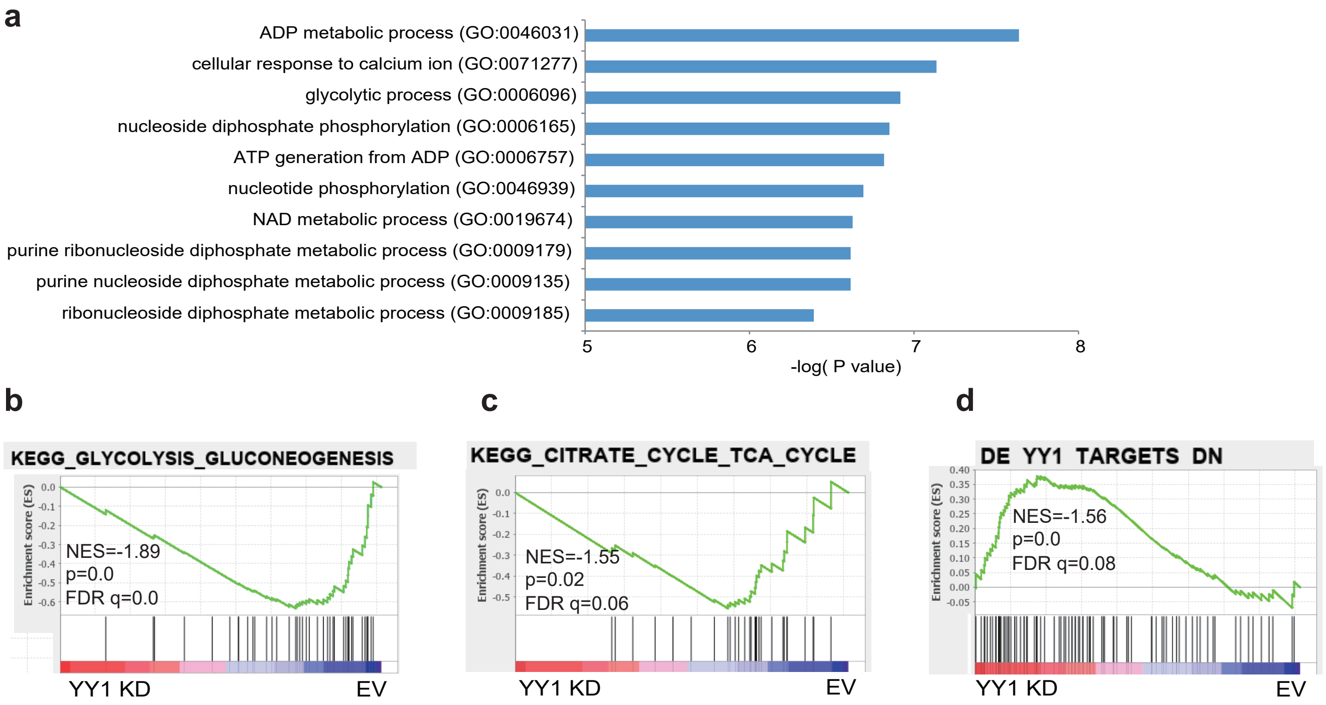


**e**

| | Total# of raw reads | Total# of raw reads after alignment |
|---|---|---|
| 22Rv1_(DHT-)_YY1_ChIP | 53367756 | 51516769 |
| 22Rv1_(DHT+)_YY1_ChIP | 52740569 | 50658309 |

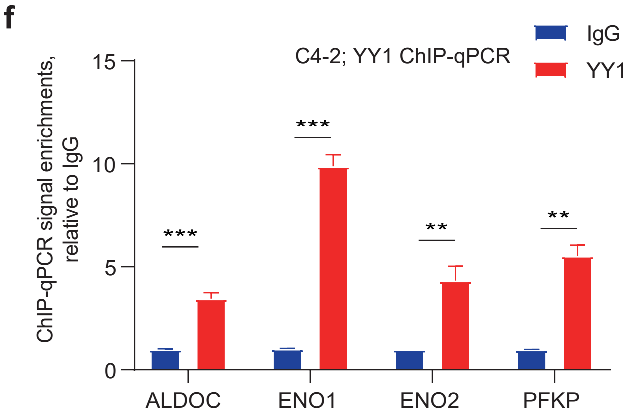


**Supplementary Fig.3: YY1 directly binds and positively regulates the metabolic genes in prostate tumor. (a)** Gene ontology (GO) analysis revealing enrichment of the indicated gene pathways in the down-regulated genes due to depletion of YY1 in the C4-2 cells, relative to EV controls. **(b-d)** GSEA revealing that, relative to mock, YY1 KD is positively correlated with downregulation of the indicated gene sets related to glycolysis **(b)** or TCA cycle **(c)**, and negatively correlated with upregulation of the indicated YY1-repressed gene set **(d)** in C4-2 cells. **(e)** Summary of the raw and uniquely aligned read counts in the YY1 ChIP-seq dataset obtained from in the vehicle- (DHT-) or dihydrotestosterone-treated (DHT+) 22Rv1 cells. **(f)** ChIP-qPCR of YY1 binding at promotor of the indicated glycolytic gene in C4-2 cells. Y-axis shows averaged fold-change ±SD of three independent experiments after normalization to inputs and then to IgG control signals.

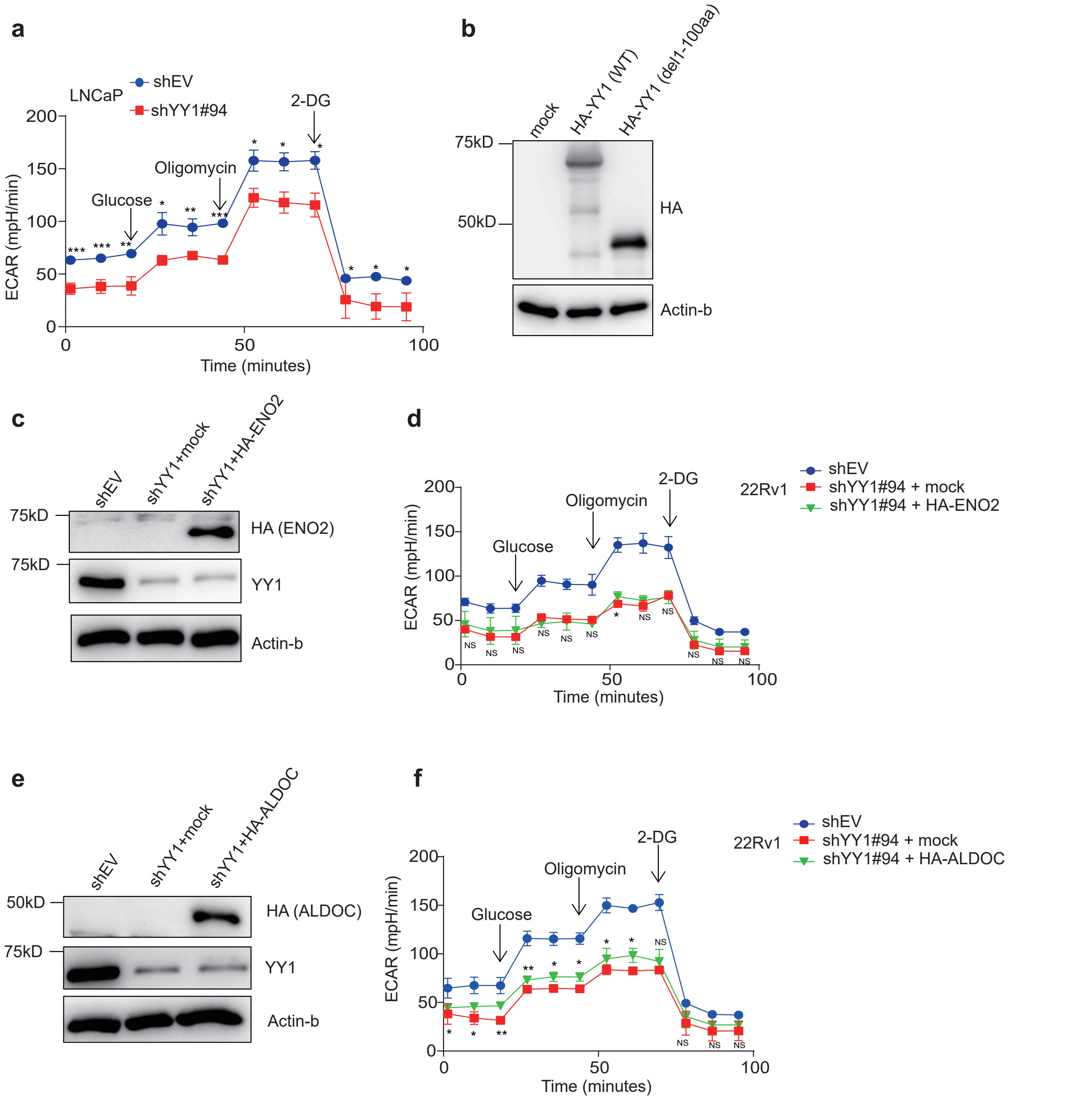


**Supplementary Fig.4: YY1 potentiates prostate tumor cell glycolysis via PFKP.(a)** Measurement of extra cellular acidification rate (ECAR) in LNCaP post-KD of YY1, compared to mock (shEV), by using the Seahorse XF 24 extracellular flux analyzer. Injection of compounds during the assay is highlighted. Data in y-axis are shown as mean ± SEM. *P<0.05;**P<0.01,***p<0.001.**(b)** HA immunoblots using the C4-2 cells post-transduction of empty vector or the indicated HA-tagged YY1, either WT or with a N-terminal transactivation domain deleted (Δ1-100aa). **(c-d)** Endogenous YY1 and anti-HA immunoblotting (panel **c**) and ECAR measurements **(d)** post-transduction of an HA-tagged ENO2 (**c**, lane 3), relative to empty vector mock (**c**, lane 2), into the YY1-depleted 22Rv1 cells (lanes 2-3). The shEV-transduced cells (lane 1) serve as control. Data shown in y-axis of panel **d** are shown as mean ± SEM. **(e-f)** Endogenous YY1 and anti-HA immunoblotting (panel e) and ECAR measurements **(f)** post-transduction of an HA-tagged ALDOC (**e**, lane 3), relative to empty vector mock (**e**, lane 2), into the YY1-depleted 22Rv1 cells (lanes 2-3). The shEV-transduced cells (lane 1) serve as control.

**PFKP-promoter(-575-+37)**

AACTGCGGGGTTTCCACCCGCCCCGGCGCTCGCGCCCACTCACGGTCTCCCTCGCCC
CCCCGGGAGATGCTCCCGGCGTTCTATCCCGCCCCGCCCTGGCGCGCCCCGCAAACG
ATGTGACCCCGGCCCCCTGCGGTCTCCCCTCCTCCGGACGCTGCCCCTGCCCCACCTG
TGGCGCTCCCGTCATCTCTAGAGCCCCCAACCAGAGTCCTCCAGCCCACCCGCGAGG
CTCGCGCACCTGCGATGCGGCCCCACTAGCAGGGACCCGCCCACCTGTGGCGCTACC
CCTGCCTCTGCCGCCTCCCGCACCGCTAGCGCGGCCCCCACCTACGACGACCCCCGC
GTAGATGCTCCTGCCCACACCCGCCCCAGCCCCGGCCCCTGCCCCTAGCCGCGCCCG
CGCCCCCTCCCCCGCAGCCCCTCCCATACGCTCGGGCCCCGCCCGTCACCGCCCATTG
GCGCTGGGGCCGGGCGGGGGCGCGGGCGGGGCGGCGGTTCCGAGTCAGGCGCG
CGCGGGCAGGGTCCCCATTGCCTGCTGCGCACCCGGACGTGCGGCTCCCCTCGGCC
TCCTCGCCATGGACGCGGACGACTCCCGGGCCCCCAAGGGCTCCT
+1

**Supplementary Fig.5: PFKP is a direct onco-target of YY1 in prostate cancer.** The nucleotide sequence of the PFKP promoter (-575 to +37), which contains four putative YY1 core binding sites as highlighted in red.

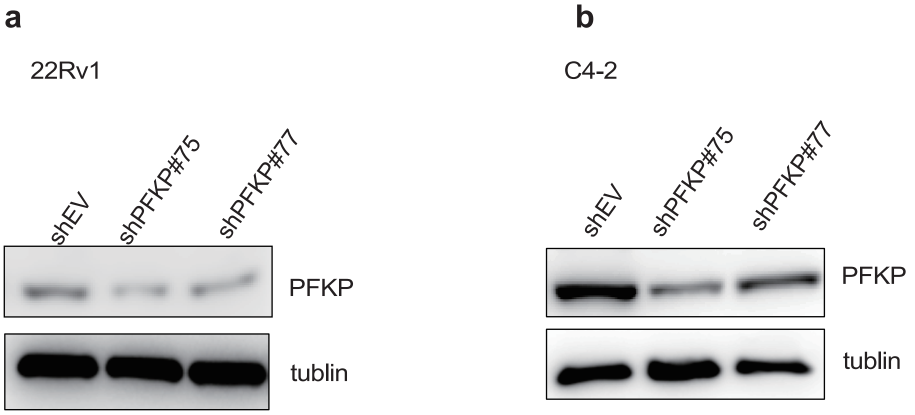


**Supplementary Fig.6: PFKP is critically involved in prostate cancer tumorigenesis in vitro and in vivo.(a-b)** Immunoblotting for PFKP in 22Rv1 **(a)** and C4-2 cells **(b)** after PFKP depletion (sh#75 or sh#77), compared to control (shEV).

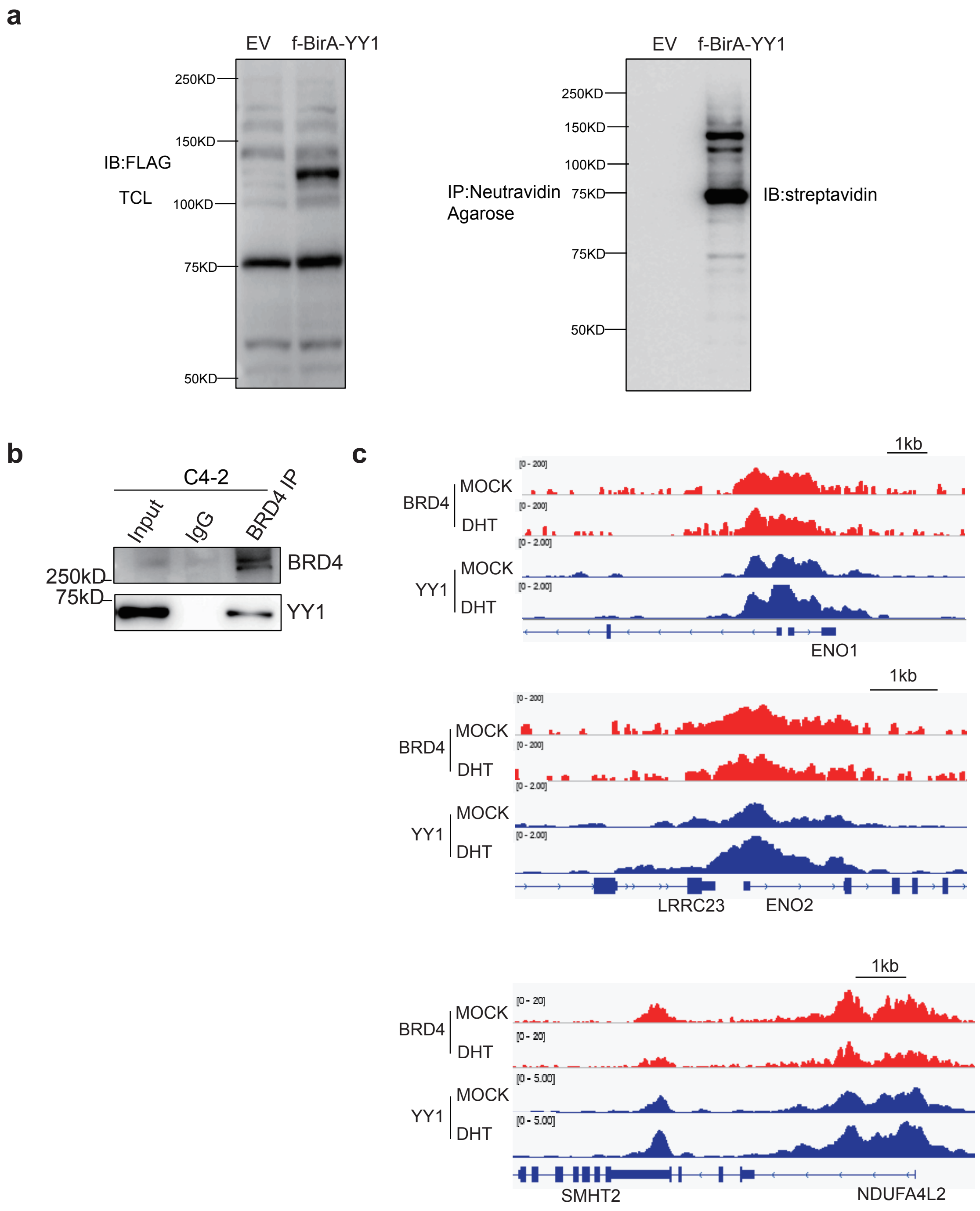


**Supplementary Fig 7. BRD4 acts as co-activator of YY1, potentiating expression of glycolysis-related genes in prostate cancer cells.**
(a) Blots for the BirA-YY1 fusion in the 22Rv1 cells stably expressing either empty vector with BirA only control (EV) or a Flag-tagged BirA-YY1 (left panel; anti-Flag) and for the biotinylated proteins (right; probed with streptavidin-HRP conjugate) post-treatment of the two cells with 50uM of biotin for 24hrs.
(b) CoIP between endogenous BRD4 and YY1 in C4-2 cells.
(c) IGV views of the YY1 and BRD4 ChIP-seq profiles at the indicated glycolytic genes in either vehicle- (MOCK) or DHT-treated (DHT+) 22Rv1 cells.

**Supplementary Table 1. RNA-seq identifies differentially expressed genes (DEGs) showing significant downregulation in the 22Rv1 cells with YY1 knockdown (shYY1), in comparison to control(shEV).**
DEGs are defined with a cut-off of padj less than 0.01 and log2|FoldChange| over 0.58 for transcripts with mean values of at least 10.

**Supplementary Table 2. RNA-seq identifies differentially expressed genes (DEGs) showing significant downregulation in the C4-2 cells with YY1 knockdown (shYY1), in comparison to control(shEV).**
DEGs are defined with a cut-off of padj less than 0.01 and log2|FoldChange| over 0.58 for transcripts with mean values of at least 10.

**Supplementary Table S3. List of the YY1 signature genes, as defined by the significant down-regulation after knockdown of YY1, relative to mock, in both 22Rv1 and C4-2 CRPC cells.**

DEGs are defined with a cut-off of padj less than 0.01 and log2|FoldChange| over 0.58 for transcripts with mean values of at least 10.

| | | | | |
|---|---|---|---|---|
| ABHD1 | DMKN | ITM2C | PHF19 | TMEM151A |
| ADAM11 | DNAAF3 | KCNMA1 | PHF21B | TMTC1 |
| ADRBK2 | DNAH2 | KCNMB4 | PLA2G12A | TNNT1 |
| ADSSL1 | DOCK6 | KIAA1467 | PNMA2 | TNS1 |
| AK7 | DPYSL2 | KIF1A | PPAP2C | TPBGL |
| ALDOC | ECE1 | L1CAM | PRR15 | TSPAN9 |
| AP3B2 | EEF1A2 | LINGO3 | PTK7 | TTC5 |
| APBB1 | EFNB3 | LPAR3 | PYGL | TUBA1A |
| APLN | EIF4E3 | LRP1 | RAB39B | TUBB4A |
| ARHGDIG | EML2 | LTBP2 | RASGRP2 | VIM |
| ARHGEF17 | EPOR | LTBP4 | RASSF5 | WNT7B |
| ARHGEF2 | ERCC2 | MAPT | REEP6 | **YY1** |
| ARHGEF25 | FANCE | MAST1 | RNF150 | ZSCAN18 |
| ASIC1 | FBLN1 | MATK | RNF157 | |
| ASMTL | FCGRT | MCAM | S100A13 | |
| ATP1A3 | FSTL3 | MOCS1 | S100P | |
| BEX1 | GALNT12 | MSI1 | SALL2 | |
| CACNA2D2 | GLDC | NAP1L2 | SCD | |
| CAMK2B | GNG12 | NDUFA4L2 | SCN8A | |
| CDH23 | GPR27 | NGFRAP1 | SCRN1 | |
| CDK5R2 | GPX8 | NRM | SEMA3G | |
| CELSR1 | GSTM3 | NXPH4 | SLC22A17 | |
| CELSR3 | H19 | OBSL1 | SLC6A8 | |
| CERCAM | H2AFY2 | OGDHL | SLIT1 | |
| CHCHD10 | HMGCS2 | PALM | SOBP | |
| CHGB | HOXA7 | PBX1 | SORBS2 | |
| CHMP1B | HTRA1 | PCYOX1L | SORL1 | |
| COLGALT2 | IFITM2 | PDZD4 | SPEG | |
| D4S234E | IFITM3 | PEPD | SYT17 | |
| DAAM1 | INSIG1 | **PFKP** | TCEA2 | |

**Table S4. Information of the used primers.**

Primers used in qRT-PCR assays

| | |
|---|---|
| ACTIN | Fw:5'AGAGCTACGAGCTGCCTGAC3'<br>Rev:5'AGCACTGTGTTGGCGTACAG3' |
| YY1 | Fw:5'TGTGGCAAAGCTTTTGTTGA3'<br>Rev:5'CGCAAATTGAAGTCCAGTGA3' |
| PFKP | Fw:5'ACCTCCACAGGCACAGGATTC3'<br>Rev:5'TCAATAATACGTGCATAAAGTG3' |
| PFKL | Fw:5'GGCATTTATGTGGGTGCCAAAGTC3'<br>Rev:5'CAGTTGGCCTGCTTGATGTTCTCA3' |
| ENO1 | Fw:5'GCTCTCCCCAGGCGTTCAATGTC3'<br>Rev:5'CTTCTTTATTCTCCAGGATG3' |
| ENO2 | Fw:5'TCTCTGCAGTGGTCTCCATTG3'<br>Rev:5'GCTTGGATGGCTTCAGTGAC3' |
| ALDOC | Fw:5'CCCCTAGGCAGCATGGCCAAGCG3'<br>Rev:5'CTTGATGCCCACGACGATGCCC3' |
| NDUFA4L2 | Fw:5'CTGGGACAGAAAGAACAACCCG3'<br>Rev:5'CAGCCTGGCTTAGAAGTCTGGC3' |
| OGDHL | Fw:5'GTACGCAGACAAGCTGATTGCC3'<br>Rev:5'ATCCTTGGACCTGCCATAAGCC3' |

Primers used in CHIP-PCR assays

| | |
|---|---|
| ALDOC | Fw:5'GATCCGCAAACAGATGAGG3'<br>Rev:5'GGAGGAGTCACGTAGCTCTG3' |
| ENO1 | Fw:5'TCGTGGCCATCTCCACTATC3'<br>Rev:5'GTTTTTGGGATAGACGCAGA3' |
| ENO2 | Fw:5'TGCAGCTGCGCACCTGCTCC3'<br>Rev:5'AGAGGGCCAGAAAGCGAAGG3' |
| PFKP | Fw:5'TTTCCGCTGGGAGATTTTTA3'<br>Rev:5'5'TGGTTTTCGTTTTTGTCACG3' |

**Supplementary Materials and Methods**

**Immunohistochemistry (IHC).** YY1 IHC staining was conducted using a rabbit anti-YY1 antibody (Atlas Antibodies #HPA001119) and the Bond fully automated slide staining system (Leica Microsystems). Slides were dewaxed in Bond Dewax solution (AR9222) and hydrated in Bond Wash solution (AR9590). Heat-induced antigen retrieval was performed for 30 min at 100ºC in Bond-Epitope Retrieval solution 1 pH-6.0 (AR9961), followed with a 5-min Bond peroxide blocking step (DS9800). After pretreatment, slides were incubated for 30mins with primary antibody YY1(1:150) followed with Vector Immpress HRP anti-rabbit IgG (MP-7401-15). Chromogenic detection of all antibodies was performed using the Bond Intense R Detection kit (DS9263). Stained slides were dehydrated and covered with a slip. Positive and negative controls (no primary antibody) were included for each run. Stained slides were digitally scanned at 20x magnification using Aperio ScanScope-XT (Aperio Technologies, Vista, CA). The images were uploaded to the Aperio eSlideManager database (Leica Biosystems Inc; eSlideManager version 12.3.3.7075) at the Translational Pathology Laboratory at UNC.

**Cell Lines.** HEK293 and HEK293T cells (acquired from American Type Culture Collection, ATCC) were cultured in DMEM supplemented with 10% FBS and 1% antibiotics. The human prostate cancer cell lines, 22Rv1, C4-2 and LNCaP, were obtained from ATCC and grown in the RPMI-1640 base medium supplemented with 10% FBS and 1% antibiotics. For compound treatment experiments, cells are first cultured under ligand-starved conditions for three days using the phenol red-free RPMI-1640 base medium supplemented with 10% charcoal-stripped serum, followed by treatment with vehicle, dihydrotestosterone (DHT), or DHT together with compounds. All cells were maintained at 37 °C with 5% CO2. Authentication of cell line identities, including those of parental and derived lines, was ensured by the Tissue Culture Facility affiliated to UNC Lineberger Comprehensive Cancer Center with the genetic signature profiling and fingerprinting analysis. Every 1–2 months, a routine examination of cell lines in culture for any possible mycoplasma contamination was performed using commercially available detection kits (Lonza).

**Antibodies.** Antibodies used in the work included the rabbit antibodies against HA tag (Cell Signaling Technology C29F4), PFKP (Cell Signaling Technology 5412S), actin-beta (Cell Signaling Technology 13E5), BRD4 (Bethyl Laboratories A301-985A100), as well as the mouse antibodies against Flag tag (Sigma F1804), YY1 (Santa Cruz SC7341; for blotting), YY1 (Atlas Antibodies #HPA001119; for IHC) and α-Tubulin (Sigma T9026). Anti-FLAG M2 Magnetic Beads (F1804) was obtained from Sigma. Anti-mouse IgG (7076S) and anti-rabbit IgG (7074S) HRP-linked secondary antibodies were obtained from Cell Signaling Technology.

**Chemicals.** Dihydrotestosterone (DHT) is purchased from Sigma and the bromodomain inhibitor JQ1 were described before[1].

**Plasmids.** cDNAs of human YY1, PFKP, ENO2 and ALDOC were amplified by PCR using total RNAs from 22Rv1 cells, fused in-frame with a HA tag and then cloned into a lentiviral vector of pCDH-EF1α-MCS-IRES-Puro/Neo (System Biosciences, CD532A-2/CD533A-2). The YY1 mutations, S365D and a N-terminal deletion of Δ1-100aa, were produced by site-directed mutagenesis (Strategene) and PCR, respectively. A PGL3 luciferase reporter with a PFKP promoter (-2008 to +7) region was kindly provided by Dr. Kyung-Sup Kim. The PFKP promoter (-575 to +37) region was cloned by PCR from using the following primers: 5'-AACTGCGGGGTTTCCACCCGCCCCG-3' and 5'-AGGAGCCCTTGGGGGCCCGGGAGTC-3', followed by cloning into pGL3-basic vector (Promega). The PFKP promoter (-575 to +37) with the YY1-motif mutants were produced by site-directed mutagenesis (with the YY1 core motif sequence 5'-CCAT mutated to 5'-CGGT). The full-length wild-type BRD4 and ΔBD domain mutant were amplified from cDNAs of 22Rv1 cells and subcloned into a home-made pcDNA-3.1_Flag_tag vector for transient expression. All plasmid sequences were verified by sequencing. For the BioID-based protein interactome study, a biotin ligase (BirA) cDNA (a kind gift of BD Strahl) was cloned into the home-made MSCV-puro retroviral vector and then a Flag-tagged YY1 cDNA was fused in-frame to C-terminus of BirA. All plasmids were confirmed by sequencing before use.

**Co-immunoprecipitation (Co-IP).** Briefly, the cell pellets were lysed in RIPA buffer supplemented with complete protease inhibitors (Roche) and PMSF. 1 mg protein from whole cell lysate was incubated with the indicated antibodies on a rotator overnight at 4℃°. Then, 20 uL of protein G agarose beads (Roche, 11243233001) were added for an additional 2 h with rotation at 4℃°. The bound protein complexes were washed with RIPA buffer 3 times and then resuspended in 40ml of 2 X protein loading buffer, and boiled at 90 degree for 5 min before loading onto SDS-PAGE gel. Western blot was performed with standard protocols and PVDF membrane, and signals visualized with an ECL system as described by the manufacturer (GE healthcare).

**CRISPR/cas9-mediated gene knockout and transient RNA interference.** SgRNAs targeting YY1 were cloned into a pLenti LRG-2.1_Neo vector (Addgene 125593). A lentiviral plasmid that allows doxycycline-inducible expression of SpCas9 was obtained from Dr. David Sabatini. The sgRNA target sequences for YY1 are as follows (YY1sg1: GTGGGCGGCGACGACTCGGA; YY1sg2: GTCGGGTCGTCGGTGACCAG). Lentiviral pLKO.1-based shRNA vectors for knockdown (KD) of human YY1 (TRCN0000019894 and TRCN0000019898) or PFKP (TRCN0000037775 and TRCN0000037777) were purchased from Sigma. All plasmid sequences were verified by sequencing. YY1 ON-TARGETplus SMART-pool siRNA (L-011796-00-0005) and control siRNA (D-001810-10-05) were purchased from Dharmacon RNAi Technologies.

**Virus Production and stable cell line generation.** Generation of stable knockdown lines using the pLKO.1 lentiviral shRNA-expressing system was carried out according to providers' protocols as described before [2]. The shRNA plasmids and packaging vectors (VSV-G and psPAX2) were co-transfected using Lipofectamine 3000 into 293FT cells. Viral supernatant was collected and filtered 48 h after transfection. Cells were infected for 48 h with virus in the presence of 8 μg/ml polybrene. Then, the target cell lines were selected with the appropriate antibiotics.

**Cell proliferation assays.** 3,000 cells per well were seeded in triplicate in 96-well plates for each time point. The change in cell number was measured using MTT assay kit based on instructions of the manufacturer (Promega).

**Colony formation assays.** Cells were plated in triplicate at a density of 20,000 cells per well of 6-well plates and grown for 3 weeks before staining with iodonitrotetrazoliuim chloride solution (sigma). Fresh medium was changed twice a week.

**Luciferase reporter assays.** Cells were seeded in 24-well plates and transfected with the indicated plasmids. pRL-CMV Renilla was used as an internal control. Luciferase activity was measured 48 h after transfection using the Dual-luciferase Reporter Assay

System (Promega). Data were normalized to Renilla luciferase. Luciferase data are reported as mean ± s.e.m. of three independent experiments performed in triplicate.

**Real-time RT-PCR.** Total RNA was isolated using the RNeasy Mini Kit (Qiagen). First strand cDNA was synthesized using the High-Capacity cDNA Reverse Transcription Kit (Applied Biosystems). Real-time PCR was performed in triplicate using the iTaq Universal SYBR Green Supermix (Bio-Rad) and the Quant Studio 6 Flex Real-Time PCR System (Applied Biosystems). All values were normalized to beta-actin levels. Information of the used primers is listed in Supplementary Table 2.

**RNA-seq and data analysis.** The fastq files were aligned to the GRCh38 human genome (GRCh38.d1.vd1.fa) using STAR v2.4.2[3] with the following parameters: --outSAMtype BAM Unsorted --quantMode TranscriptomeSAM. Transcript abundance for each sample was estimated with salmon v0.1.19 [4] to quantify the transcriptome defined by Gencode v22. Gene level counts were summed across isoforms and genes with low counts (maximum expression < 10) were filtered for the downstream analyses. We tested genes for differential expression in DESeq2 [5] in R. Genes with | log2FC (fold-change) | > 0.58 and adjusted p value < 0.01 between YY1 knockdown and vector control samples were called as differentially expressed genes (DEGs).

**ChIP-seq and data analysis.** ChIP-seq reads were aligned to the human reference genome (hg19) by the BWA (V0.7.12; default parameters) software [6]. After duplicated reads were removed, MACS2 (v2.1.0; -q 0.1 -m 20 100) [7] was used to call peaks with input as controls. Peaks overlapping (>= 1 bp) with the ‘‘blacklist’’ regions identified by the ENCODE project were removed. The filtered peaks were assigned to annotated (coding and non-coding) genes and defined sequentially as ‘‘promoter’’ (±2kb of transcription start site, TSS), within “gene body”, ‘‘distal,’’ (i.e., ‘‘enhancer’’; -50kb to -2kb of TSS or +2kb of TSS to +5kb of transcription ends), or otherwise ‘‘intergenic’’ using the human RefSeq annotation. The ChIP-seq read densities were calculated and visualized as heatmaps using the program seqMINER [8]. The enrichment of motifs was identified by the software HOMER [9] with default parameters.

**GSEA.** Gene set enrichment analysis (GSEA) was performed as previously described [10] with the downloaded GSEA software (www.broadinstitute.org/gsea) by exploring the Molecular Signatures Database (www.broadinstitute.org/gsea/msigdb/annotate.jsp).

**Seahorse assay.** 22Rv1 and C4-2 cells were first cultured in RPMI-1640 medium without phenol red for 3 days. Then, the cells were harvested, counted, resuspended in assay medium (Agilent, 103576-100) and plated in XF24 assay plates (Agilent, 100777-004) at a density of 1 x 105 cells per well. The cells were kept at 37°C without CO2 for 1 h. Oxygen consumption rate (OCR) was measured using Cell Mito Stress Kit (Agilent, 103015-100). Mitochondria inhibitors were sequentially injected at the following concentrations: 1.5 μM oligomycin; 1 μM FCCP and 0.5 uM Antimycin A/ Rotenone. Extracellular acidification rate (ECAR) was measured using Glycolysis Stress Kit (Agilent, 103020-100). The cells were metabolically perturbed by sequential injections of 10mM glucose, 1 uM oligomycin and 50 mM 2-deoxyglucose. OCR and ECAR levels were recorded using a Seahorse XF 24 extracellular flux analyzer following manufacturer's instructions (Seahorse Biosciences, North Billerica, MA).

**Bio-ID.** A proximity labeling-based BioID was carried out as described[11,12]. In brief, the 22Rv1 cells that stably express a flag-tagged BirA ligase fused in-frame to the N-terminus of YY1 were treated with 50uM of biotin for 24hrs, followed by lysis in the RIPA buffer (10%

glycerol, 25mM Tris-HCl pH 8, 150mM NaCl, 2mM EDTA, 0.1% SDS, 1% NP-40, 0.2% Sodium Deoxycholate) freshly supplemented with 1X protease inhibitor cocktail, 1 mM PMSF and 250 units of Benzonase for 1h with rotation at 4 ºC. Cells stably expressing a vector with the BirA ligase only (empty vector or EV) were used as a negative control for BioID. Samples were snap frozen on liquid nitrogen and thawed on ice, and then transferred to 1.7ml Eppendorf tubes. After brief vortex of cell lysate and centrifugation at top speed for 30 minutes at 4 ºC, the cleared supernatant was collected and incubated with the Neutravidin beads (Thermo Fisher, 29204) overnight at 4 ºC. The beads were washed twice with the RIPA buffer (with no additives), twice with the TAP lysis buffer (10% glycerol, 150mM NaCl, 2mM EDTA, 0.1% NP-40, 50mM HEPES pH 8) and three times with the ABC buffer (50mM Ammonium bicarbonate, pH 8), followed by spin at 400g for 2 min at 4 ºC. The bead:protein samples can be stored in 100ul of the ABC buffer before mass spectrometry-based protein identification.

**Mass Spectrometry-based Protein Identification and Proteomic Data Analysis**

Proteins bound on beads were eluted by adding 100 µL of 1x Laemmli buffer (Boston Bioproducts) and heating at 95 degrees for 5 minutes. Supernatants were then loaded on a 4-12% Bis-Tris Deep Well gel, resolved by one-dimensional SDS-PAGE and visualized by Coomassie staining. Each SDS-PAGE gel lane was sectioned into 12 segments of equal volume. Each segment was subjected to in-gel trypsin digestion by using the following established protocol. Gel slices were de-stained in 50% methanol (Fisher), 50 mM ammonium bicarbonate (Sigma-Aldrich), followed by reduction in 10 mM Tris(2-carboxyethyl)phosphine (TCEP; Pierce) and alkylation in 50 mM iodoacetamide (Sigma-Aldrich). Gel slices were then dehydrated in acetonitrile (Fisher), followed by addition of 100 ng porcine sequencing grade modified trypsin (Promega) in 50 mM ammonium bicarbonate (Sigma-Aldrich) and incubation at 37 degrees for 12-16 hours. Peptide products were then acidified in 0.1% formic acid (Pierce). Tryptic peptides were separated by reverse phase XSelect CSH C18 2.5 um resin (Waters) on an in-line 150 x 0.075 mm column using a nanoAcquity UPLC system (Waters). Peptides were eluted using a 30 min gradient from 97:3 to 67:33 buffer A:B ratio (Buffer A = 0.1% formic acid, 0.5% acetonitrile; buffer B = 0.1% formic acid, 99.9% acetonitrile). Eluted peptides were ionized by electrospray (2.15 kV) followed by MS/MS analysis using higher-energy collisional dissociation (HCD) on an Orbitrap Fusion Tribrid mass spectrometer (Thermo) in top-speed data-dependent mode. MS data were acquired using the FTMS analyzer in profile mode at a resolution of 240,000 over a range of 375 to 1500 m/z. Following HCD activation, MS/MS data were acquired using the ion trap analyzer in centroid mode and normal mass range with precursor mass-dependent normalized collision energy between 28.0 and 31.0.

Proteins were identified by searching the UniProtKB database using Mascot (Matrix Science), Scaffold (Proteome Software) was used to verify MS/MS based peptide and protein identifications. Peptide identifications were accepted if they could be established with less than 1.0% false discovery by the Scaffold Local FDR algorithm. Protein identifications were accepted if they could be established with less than 1.0% false discovery and contained at least 2 identified peptides. and the counts were normalized to log2 normalized spectral abundance factor (NSAF) values. Significant interacting proteins were identified by a log2 fold change >1.

**Reference used in Supplementary:**


1 Filippakopoulos, P. *et al.* Selective inhibition of BET bromodomains. *Nature* **468**, 1067-1073, doi:10.1038/nature09504 (2010).

2 Cai, L. *et al.* ZFX Mediates Non-canonical Oncogenic Functions of the Androgen Receptor Splice Variant 7 in Castrate-Resistant Prostate Cancer. *Mol Cell* **72**, 341-+, doi:10.1016/j.molcel.2018.08.029 (2018).

3 Dobin, A. *et al.* STAR: ultrafast universal RNA-seq aligner. *Bioinformatics* **29**, 15-21, doi:10.1093/bioinformatics/bts635 (2013).

4 Patro, R., Duggal, G., Love, M. I., Irizarry, R. A. & Kingsford, C. Salmon provides fast and bias-aware quantification of transcript expression. *Nat Methods* **14**, 417-419, doi:10.1038/nmeth.4197 (2017).

5 Love, M. I., Huber, W. & Anders, S. Moderated estimation of fold change and dispersion for RNA-seq data with DESeq2. *Genome Biol* **15**, 550, doi:10.1186/s13059-014-0550-8 (2014).

6 Li, H. & Durbin, R. Fast and accurate long-read alignment with Burrows-Wheeler transform. *Bioinformatics* **26**, 589-595, doi:10.1093/bioinformatics/btp698 (2010).

7 Zhang, Y. *et al.* Model-based Analysis of ChIP-Seq (MACS). *Genome Biol* **9**, doi:ARTN R137
10.1186/gb-2008-9-9-r137 (2008).

8 Ye, T. *et al.* seqMINER: an integrated ChIP-seq data interpretation platform. *Nucleic Acids Res* **39**, doi:ARTN e35
10.1093/nar/gkq1287 (2011).

9 Heinz, S. *et al.* Simple Combinations of Lineage-Determining Transcription Factors Prime cis-Regulatory Elements Required for Macrophage and B Cell Identities. *Mol Cell* **38**, 576-589, doi:10.1016/j.molcel.2010.05.004 (2010).

10 Xu, B. W. *et al.* Selective inhibition of EZH2 and EZH1 enzymatic activity by a small molecule suppresses MLL-rearranged leukemia. *Blood* **125**, 346-357, doi:10.1182/blood-2014-06-581082 (2015).

11 Roux, K. J., Kim, D. I., Burke, B. & May, D. G. BioID: A Screen for Protein-Protein Interactions. *Curr Protoc Protein Sci* **91**, 19 23 11-19 23 15, doi:10.1002/cpps.51 (2018).

12 Roux, K. J., Kim, D. I. & Burke, B. BioID: a screen for protein-protein interactions. *Curr Protoc Protein Sci* **74**, 19 23 11-19 23 14, doi:10.1002/0471140864.ps1923s74 (2013).
